# Spatially-extended nucleation-aggregation-fragmentation models for the dynamics of prion-like neurodegenerative protein-spreading in the brain and its connectome

**DOI:** 10.1101/692038

**Authors:** Sveva Fornari, Amelie Schäfer, Ellen Kuhl, Alain Goriely

## Abstract

The prion-like hypothesis of neurodegenerative diseases states that the accumulation of misfolded proteins in the form of aggregates is responsible for tissue death and its associated neurodegenerative pathology and cognitive decline. Some disease-specific misfolded proteins can interact with healthy proteins to form long chains that are transported through the brain along axonal pathways. Since aggregates of different sizes have different transport properties and toxicity, it is important to follow independently their evolution in space and time. Here, we model the spreading and propagation of aggregates of misfolded proteins in the brain using the general Smoluchowski theory of nucleation, aggregation, and fragmentation. The transport processes considered here are either anisotropic diffusion along axonal bundles or discrete Laplacian transport along a network. In particular, we model the spreading and aggregation of both amyloid-*β* and *τ* molecules in the brain connectome. We show that these two models lead to different size distributions and different propagation along the network. A detailed analysis of these two models reveals the existence of four different stages with different dynamics and invasive properties.

## 1 Introduction

Neurodegenerative diseases such as Alzheimer’s (AD) or Parkinson’s (PD) are devastating conditions associated with a systematic destruction of brain tissues leading to cognitive decline, neurobehavioral symptoms, and eventually death. While for PD there exist some treatments to alleviate some of the symptoms, there is no known cure for any of these diseases. Post-mortem analyses of brain tissues affected by neurodegenerative diseases reveal the presence of protein aggregates. For instance, in the case of AD, extracellular amyloid-beta (A*β*) plaques and intracellular neurofibrillary tangles of tau (*τ*) proteins are observed and correlated with the evolution of the disease [1]. The systematic mapping of these lesions either in postmortem brains obtained at various stages of the disease or by in vivo positron emission tomography imaging provides a map of the spatiotemporal evolution of the disease [2, 3]. Unlike other diseases, neurodegenerative diseases appear to follow a predictable spreading pattern through the brain. For instance, in AD, *τ* aggregates are first found in the locus coeruleus and entorhinal cortex and then evolves to the hippocampus, the temporal cortex, the parietal cortex before invading the motor cortex and occipital areas [4]. In the last stage of the disease, all cortical areas are affected and the patient condition rapidly declines. Different neurodegenerative diseases exhibit different invasion patterns associated with different initial seeding zones and specific protein aggregates.

These systematic invasion patterns of protein aggregates are the basis of the *prion-like* hypothesis for neurodegenerative diseases. This mechanism is based on the idea that, like prion diseases [5], neurodegenerative diseases are caused by the systematic aggregation and transport of misfolded proteins in the brain through the axonal pathways [6, 7, 8, 9, 10]. Specifically, it applies to *τ* protein aggregates found in AD. Tau proteins are small proteins that stabilize microtubules in the axon [11]. In healthy tissue, they are naturally produced by the cell and transported primarily along the axons where they bind to multiple microtubules. However, in some conditions, these proteins can be hyperphosphorylated and start forming misfolded aggregates. This misfolded form of the protein acts as a toxic template on which regular *τ* protein can be bound and converted to misfolded ones. These aggregates grow into increasingly larger fibrillar assemblies [7, 12] that can also fragment into smaller aggregates. Since *τ* is an intracellular protein, these various large aggregates primarily spread across the brain through the network of axonal pathways [13, 14] and various mechanisms of cell-cell spreading have been identified [15]. Similarly, it is known that A*β* form large extra-cellular aggregates. Assuming that these aggregates are transported within the brain as a simple diffusion process, we know from diffusion tensor imaging, that diffusion is preferentially along the axons. Therefore, even though these proteins are found outside the cell, they also diffuse anisotropically.

The kinetic of aggregation and fragmentation of misfolded proteins and their spatiotemporal evolution can be modeled by either following the total concentration of toxic proteins [16, 17], the concentration of healthy and toxic proteins using a heterodimer model [18], or a Smoluchowski-type model where the concentrations of polymers of different sizes are followed independently [19].

Following the size distribution is important to understand the slow time scales associated with the disease and to identify the aggregate size responsible for damage so that they can be targeted by antibodies. Therefore, we use the aggregation theory of Smoluchowski to study the spread of intracellular protein aggregates across the brain. We are particularly interested in studying to what extent coarse-grained models, which are easier to simulate, can be used to represent the complex underlying kinetics. Our approach consists in formulating the continuous problem first using anisotropic diffusion and then discretizing the equations on a network. Once the models for A*β* and *τ* propagation have been established, we analyze them in parallel to identify typical behaviors and how particular features arise from the modeling choices.

## 2 General theory of aggregation-fragmentation equations

Before we look specifically at the problem of proteins in the brain, it is of interest to consider the general theory of Smoluchowski for the aggregation and fragmentation of particles in space and time [20]. We will first consider the continuum case before discretizing these equations on a network. In this theory, we follow the concentration *c*_*i*_ of aggregates *C*_*i*_ of size *i* ∈ ℕ. The concentrations are defined both in space and time so that *c*_*i*_ = *c*_*i*_(**x**, *t*), **x** ∈ Ω ⊂ ℝ^3^, *t* ∈ ℝ. Apart from nucleation events, we consider only binary processes where the aggregates *i* and *j* interact with aggregates of size *i* + *j* with an aggregation rate *k*_*i,j*_ and fragmentation rate *β*_*i,j*_:

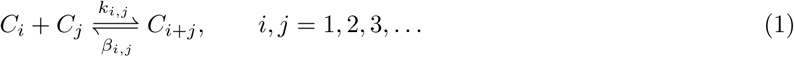

In addition, we assume that there exists a source of monomers and a process of clearance reducing each population with a constant relative rate. The general form of these equations is then

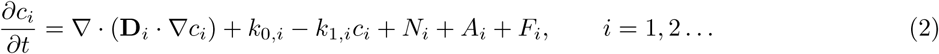

where **D**_*i*_ is the diffusion tensor characterizing the spreading of an aggregate of size *i*. We assume a source of monomers *k*_0,1_ = *γ*(**x**) and *k*_0,*i*_ = 0 for *i* > 1 and clearance terms of the *i*-mer, *k*_1,*i*_ = *k*_1,*i*_(**x**), that are possibly space-dependent. This dependence reflects the possibility that different locations may be associated with higher rates of production or clearance. The remaining terms in the equations are the nucleation term *N*_*i*_, the aggregation term *A*_*i*_, and the fragmentation term *F*_*i*_. We consider these three processes separately:

### Nucleation

We consider two different type of nucleation processes (see Fig. 1) that have shown to be important in the context of protein kinetics for neurodegenerative diseases [21, 22]. First, primary nucleation corresponds to *n*_1_ > 1 monomers forming an aggregate of size *n*_1_:

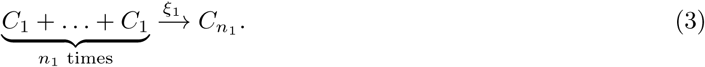

Second, we include secondary nucleations where existing aggregates facilitates the formation of new aggregates with *n*_2_ > 1 monomers [23]:

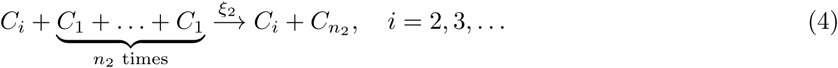

In this case the rate constant is proportional to the total mass Σ_*i*>1_ *iC*_*i*_. Taking into account both contributions, the nucleation term is given by

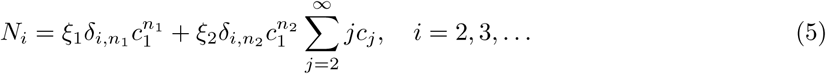

where *δ*_*i,j*_ is the usual Kronecker delta (1 when *i* equals *j* and 0 otherwise). The conservation of mass in the nucleation process implies that 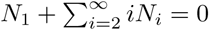. Hence, we have

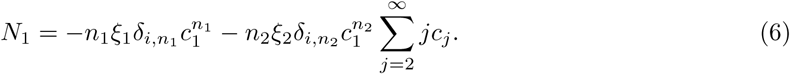

### Aggregation

Considering the possible changes in the concentration *c*_*i*_ with fixed *i* > 1, we see from (1) that the aggregate *C*_*i*_ appears in two reactions. It disappears in the presence of *C*_*j*_ to form *C*_*i*+*j*_:

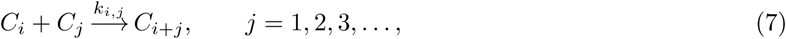

and appears in the same type of reaction but with different indices

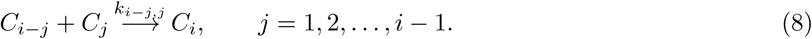

Note that the symmetry obtained by swapping *j* with *i* − *j* in this equation means that we count these reactions twice (except when *i* = 2*j*).

**Figure 1:**
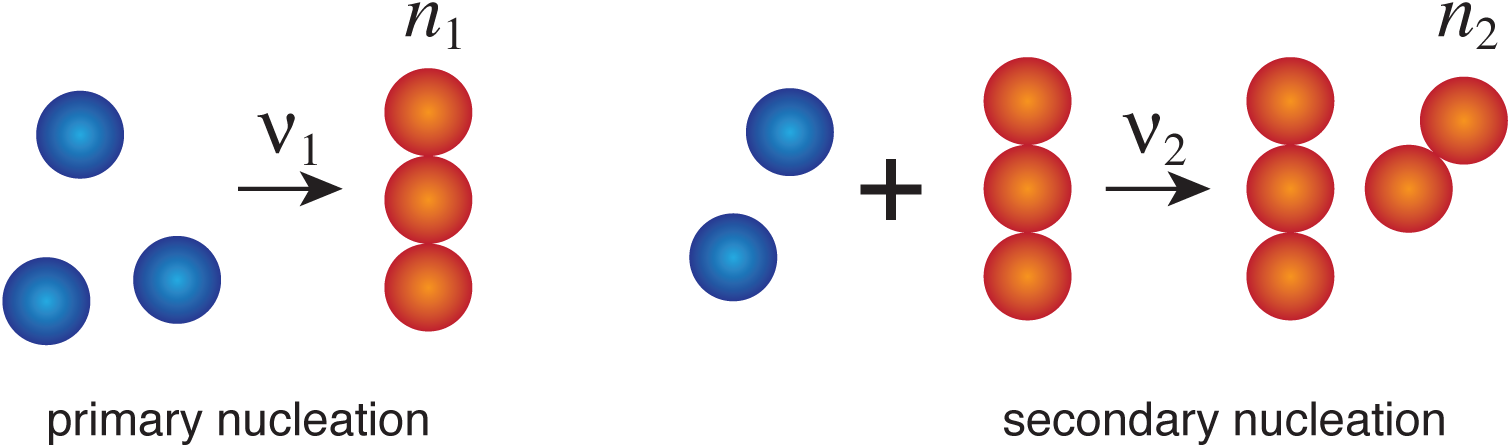
The two nucleation processes. Primary nucleation brings *n*_1_ monomers together to form an aggregate of size *n*_1_ (here *n*_1_ = 3). In secondary nucleation, the presence of an aggregate of any size, catalyzes nucleation to form an aggregate of size *n*_2_ (here *n*_2_ = 2).

Taken together these effects can be written thanks to the law of mass action as [24, 25]

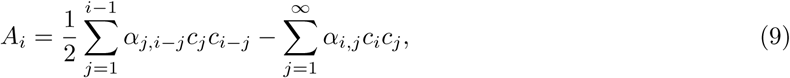

where the factor 1*/*2 appears due to the double counting and *α*_*i,j*_ = *α*_*j,i*_ = *k*_*i,j*_ when *i* ≠ *j*, and *α*_*i,i*_ = 2*k*_*i,i*_. Terms of the form 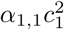 in the equation for *c*_2_ also represents the possibility of nucleation from two monomers *C*_1_ forming a dimer *C*_2_. Therefore, if the primary nucleation is binary (*n*_1_ = 2), the total nucleation rate is *α*_1,1_ + *ξ*_1_. If *n*_1_ > 2, then there is no binary nucleation and *α*_1,1_ = 0.

### Fragmentation

The fragmentation term follows the same construction and takes into account the reactions (7) and (8) in the reverse direction. We consider the loss of aggregates *C*_*i*_ by the reaction.

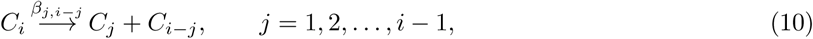

and the creation of aggregates of size *i* by the fragmentation of larger aggregates

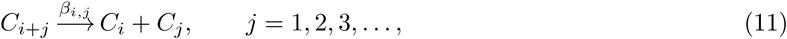

which leads to

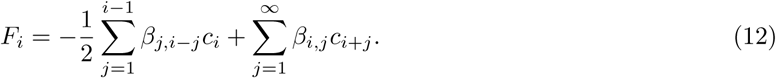

Note that since we only consider binary processes, we neglect the possibility of aggregation of more than two smaller aggregates or fragmentation processes leading to more than two aggregates of smaller sizes. If we assume that aggregation dominates fragmentation, we have *α*_*i,j*_ > *β*_*i,j*_.

Taken together, the Smoluchowski equations for nucleation-aggregation-fragmentation read

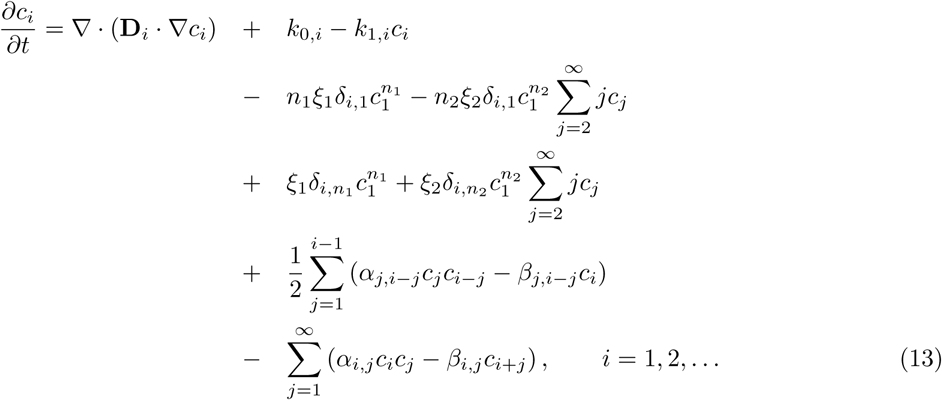

Whereas the general form of these equations is well accepted, the problem is to find the specific form of the coefficients for a given process and then solve this infinite set of nonlinear partial differential equations. If we consider aggregates of size up to *N*, the number of free parameters is of order *N*^2^. Modeling reaction rates usually rely on a combination of physical assumptions, thermodynamics, and statistical physics, all based on direct comparisons with experimental data.

## 3 Smoluchowski equations for neurodegenerative diseases

The approach discussed here has been used to study the spread of proteins in some neurodegenerative diseases (see review [26]). For Alzheimer’s disease, the emphasis in most models is on the the evolution of A*β* fibrils, which have been thought as the main responsible mechanism related to cell death. For instance, a homogeneous Smoluchowski model has been proposed by Murphy and Pallitto and validated against kinetic experiments [27, 28]. Many other homogenous models have been considered for A*β* fibrils and prion diseases. These models are obtained from (13) by taking **D**_*i*_ = **0** for all *i* and are therefore sets of ordinary differential equations. This is the classical framework of Smoluchowski equations for which there exists a large literature [29, 30, 31]. The central question is to obtain the evolution in time of the concentration distribution given general properties of the rate coefficients and whether gelation occurs. Physically, gelation refers to the process of creating very large particles in the system. Mathematically, in these equations, it corresponds to a loss of mass in the system due to a non-zero mass flux towards larger particles in the limit of particle sizes going to infinity. The main advantage of the homogenous case is that differential equations for moments of the distribution can be obtained and, in some cases, the problem can be reduced to a finite set of differential equations for these moments. This mathematical framework can then be used to fit the constants appearing in the system against experimental data [32].

Not surprisingly, the inhomogeneous case where transport is considered is much more complicated and no such moment formulation is possible. Yet, a few mathematical works have established the global existence of solutions for particular rate equations [33, 34, 35] or obtained particular solutions [36].

Of particular relevance for the present discussion is the work of Bertsch et al. who considered a model of the form (13) for the accumulation and spreading of A*β* and implemented the model in a brain slice geometry [19]. Similar equations have also been discretized and studied on networks by Matthäus who was motivated by the study of prion diseases [18, 37, 38].

Here, following the prion-like hypothesis of neurodegenerative diseases, we study and compare two different models for the aggregation and transport of either A*β* or *τ* proteins, the hallmarks of Alzheimer’s disease. According to the prion-like hypothesis, these proteins are mostly transported along axonal pathways. Hence, a network approach for the spatiotemporal evolution of these aggregates is justified. The network models are obtained as coarse-grained models of the continuum models.

It is important to make the distinction between the population of healthy proteins and the misfolded (toxic) ones. We assume that in healthy conditions, the healthy proteins have a concentration *m* = *m*(**x**, *t*). We assume that the only process by which misfolded monomers can be produced is through conversion of a healthy protein or by fragmentation of a polymer. However, our assumption about fragmentation (see below) does not allow for the loss of a single monomer. Hence, toxic monomers can only be produced by conversion of a healthy protein and will only appear in the system as they form larger fibrils. Since we pool the process of conversion and aggregation together through a single constant, there is no need to track separately the population of misfolded monomers. Hence, we use *c*_1_ = *m* as the overall population of monomers present in the system. For the dynamics to start, we must either have a nucleation mechanism with rate *κ* that describes the probability of two such monomers to come together to make a dimer of misfolded proteins, or assume that this conversion has already taken place and the system has a certain level of seeded misfolded dimers.

### 3.1 Diffusion, growth and expansion

It is important to identify the possible sources for the creation, transport, and expansion of toxic proteins. We define the total density and concentration of aggregates (excluding healthy monomers) as

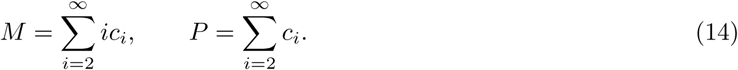

Integrated over the entire domain, these two quantities are, respectively, the total mass and total number of aggregates of toxic proteins. A typical aggregate length, measured in unit of monomer length, is obtained as the ratio of these two moments *λ*(*t*) = *M/P*. In the case of fibrils, *λ* is the mean filament length.

There are three main processes in the dynamics, each associated with its own time scale. *Diffusion* has the effect of lowering locally a high concentration by transporting aggregates in nearby regions. Hence, starting in one small region with high concentration, diffusion allows for seeds to propagate.

*Growth* refers to the evolution of the fibril length: the transfer from small aggregates to larger aggregates. This process is mostly controlled by the parameters *α*_*i,j*_ and leads to an increase of *λ*. Once a toxic seed of small size is created, growth increases the size of that seed. The process is dampened by either fragmentation or clearance.

*Expansion* refers to an increase of the total mass of toxic proteins. It is controlled by three possible sub-processes: *primary nucleation* that creates seeds directly from the pool of monomers, *secondary nucleation* that creates seeds from monomers but requires activation from other aggregates, and *fragmentation* that creates news seeds from larger aggregates at the expense of growth processes.

While the primary nucleation process is necessary to create initially toxic seeds, the two main expansions mechanisms (secondary nucleation or fragmentation) are observed for different proteins. For A*β, in vitro* experiments on the formation of oligomers based on the A*β*42 peptide have shown [39] that both primary and secondary nucleation processes are necessary to capture correctly the kinetic of the process. Once a population of toxic seed is established and grow, it acts as a catalyst for the formation of more seeds through a positive feedback mechanism. However, for *τ* proteins, primary nucleation and fragmentation is sufficient to explain homogeneous *in vitro* experiments [40]. The creation of new seeds from larger ones creates new targets for monomers to be transformed into toxic proteins and secondary nucleation is not required.

Based on these observations, we establish two classes of models. The first one for A*β* is based on primary and secondary nucleations only (no fragmentation). The second class of models, relevant for *τ* proteins, is based on primary nucleation and fragmentation only (no secondary nucleation). Both models share a number of common assumptions that we discuss now before specializing them.

### 3.2 Continuous models for fibril propagation

#### Linear aggregation

Various authors have discussed the possibility of a general aggregation mechanism from aggregates of various sizes [28, 27]. However, in the formation of neurofibrillary tangles, the growth of a fibril is dominated by the addition of monomers at the ends of the fibril. Therefore, we assume here that the main mechanism is through the formation of fibrils by the addition of monomers. This assumption considerably simplifies the equations as we only consider aggregation processes of the form

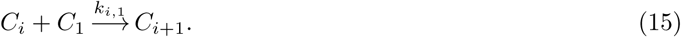

This type of coagulation kinetic is similar to the well-known Becker-Döring process that has been studied extensively [41, 42, 43, 44, 45]. The main difference is that in Becker-Döring only one monomer at most is lost during fragmentation.

We further assume that for polymers with more than two particles, the rates are independent of the size so that the probability of attaching a monomer to a chain does not depend on how long the chain is: *k*_*i*,1_ = *k*_1,*i*_ = *k* for all *i* > 2, which implies *α*_*i*,1_ = *α*_1,*i*_ = *α* for all *i* > 2. We distinguish the term involving dimers

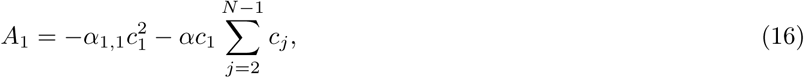

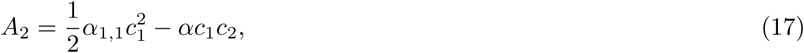

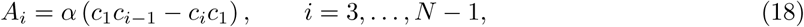

where *N* is the size of the super-particle discussed next.

#### Super-particle

Rather than considering an infinite set of equations, we consider a truncation of these equations by following the concentration of a super-particle consisting of all aggregates of size equal or larger than *N*. The value of *N* is chosen to be the size of the smallest particle that is insoluble and, therefore, does not diffuse. Hence, we have **D**_*N*_ = **0**. We further assume that the super-particle does not fragment so that *F*_*N*_ = 0. Following the argument in [19], the equation for *x*_*N*_ is

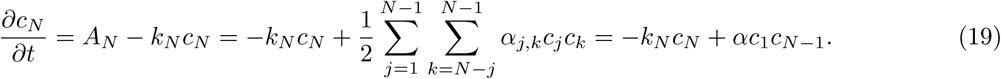

The limit *N* → ∞ recovers the classic case with infinitely many species.

#### Primary and secondary nucleation

We assume that nucleation happens through the formation of dimers (*n*_1_ = 2) as observed experimentally [46] and combine the two contribution to the creation of dimers by introducing 2*κ* = *α*_1,1_ + *ξ*_1_. When secondary nucleation takes place, we take *n*_2_ = 2 and *ξ*_2_ = *ξ*.

#### Finite fragment size

When a chain fragments, we assume that it is unlikely to lose small fragments. Hence, we assume that there is a minimal fragment size *ζ* such that fragments smaller than *ζ* cannot be produced. In a chain with *j* elements, there are *j* − 1 places where it can break. However, since small chains cannot be produced, there are only *j* – 1 − 2(*ζ* − 1) = *j* + 1 − 2*ζ* places where the chain can break. Hence, we only consider the loss of aggregates *C*_*i*_ through

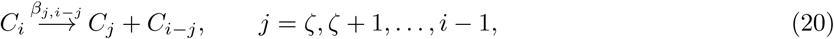

and the creation of aggregates of size *i* as

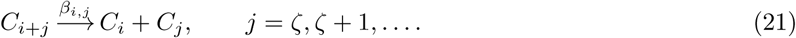

Further, since the super-particle cannot fragment, we have *F*_*i*_ = 0 if *i* < *ζ* and for *i* > *N* − *ζ* − 1. Assuming that the rate of fragmentation is independent of the size and the position at which the filament breaks, we have *β*_*i,j*_ = *β* for all *i* and *j*:

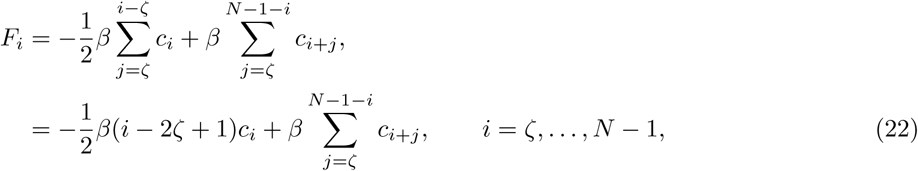

where it is understood that the sum 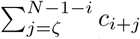 vanishes identically when the upper bound (*N* − 1 − *i*) is less than the lower bound *ζ*, which happens for *i* > *N* − *ζ* − 1. For the rest of the analysis, we will follow [40] and assume that the smallest possible fragment is of size *ζ* = 2, indicating the fact that once a dimer is formed it is stable and never fragments. For the creation of large aggregates to take place, aggregation must be favored over fragmentation, which is enforced by *β* < *α*.

#### Transport scaling

Aggregates of different sizes are not transported in the same way with larger aggregates diffusing more slowly [47]. Indeed, the diffusion coefficient of a soluble molecule scales approximately as a power of its molecular weight and the weight of an oligomer is proportional to its size. Therefore, we scale the diffusion tensor according to size by a power law of the form

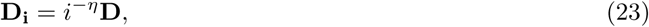

where *η* is a constant. Assuming that the diffusion constant scales inversely to the mass of the molecule, it scales as the cubic root of its length [48, 49], hence, we take *η* = 1*/*3.

For the diffusion tensor we choose [17]

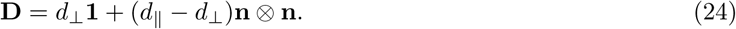

This is a transversely anisotropic diffusion tensor with a preferential diffusion *d*_‖_ (with *d*_‖_ ≫ *d*_⊥_) along the axon bundle characterized by the unit vector field **n** = **n**(**x**, *t*).

#### Clearance rate

Aggregates are continuously removed from the system through normal clearance processes such as the CSF and the glymphatic system [50]. There are two different assumptions of interest for our study.

First, we can assume that the clearance rate is independent of the size of the aggregate. In this case of *size-independent clearance*,

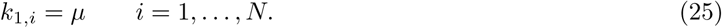

Second, we can assume that for a given phagocytic activity or antibody the clearance of an oligomer with *i*-elements is the same as the removal of each element. Therefore, chains of size *N* or larger cannot be removed and it becomes increasingly difficult to remove large chains: the *size-dependent clearance rates* are inversely proportional to the size of the oligomer:

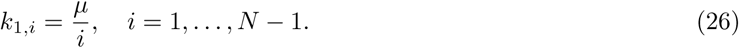

Both cases are taken into account by writing *k*_1,*i*_ = *µ*_*i*_, *i* = 1, …, *N*.

#### The continuous model

Taken into account all the above assumptions, the full equations for the concentrations take the form

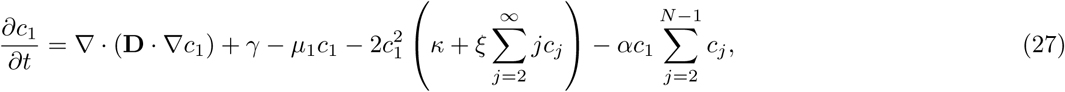

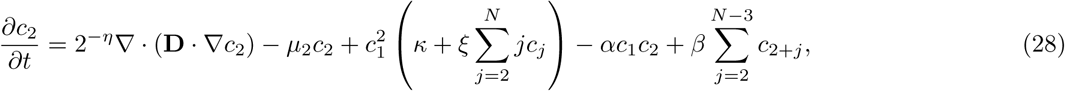

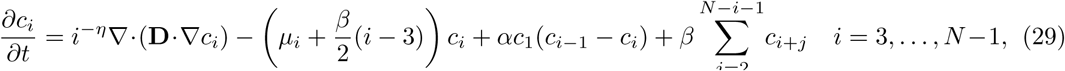

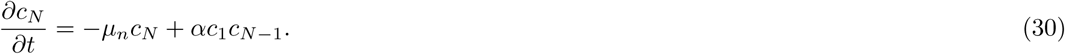

where **D** is given by (24).

The main difference between the model for A*β* and *τ*, apart from the initial seeding zones, and different sets of parameters is the choice *β* = 0 (no fragmentation) for the continuous A*β* model and *ξ* = 0 (no secondary nucleation) for the *τ* model.

### 3.3 Scaling

It is interesting to consider the respective size of the parameters and introduce a proper scaling of the parameters so that the new variables are dimensionless. In the homogeneous case and in the absence of clearance and production, the remaining parameters, given in Table 1 can be evaluated from *in vitro* experiments.

**Table 1:** Typical parameters for the *τ* model are taken from [40] and for the *β* model are from [39].

| $A\beta$ model | | | $\tau$ model | | |
| --- | --- | --- | --- | --- | --- |
| $\kappa$ | Nucleation | $6 \times 10^{-4} \text{M}^{-1} \text{s}^{-1}$ | $\kappa$ | Nucleation | $2.8 \times 10^{-4} \text{M}^{-1} \text{s}^{-1}$ |
| $\xi$ | Secondary nucleation | $1 \times 10^4 \text{M}^{-2} \text{s}^{-1}$ | $\xi$ | Secondary nucleation | 0 |
| $\beta$ | Fragmentation | 0 | $\beta$ | Fragmentation | $11.2 \times 10^{-11} \text{s}^{-1}$ |
| $\alpha$ | Elongation Rate | $6 \times 10^6 \text{M}^{-1} \text{s}^{-1}$ | $\alpha$ | Elongation Rate | $8.4 \times 10^3 \text{M}^{-1} \text{s}^{-1}$ |
| $m_0$ | Initial monomer c. | $\gamma/\mu = 10^{-6} \text{M}$ | $m_0$ | Initial monomer c. | $\gamma/\mu = 10^{-6} \text{M}$ |
| $\tilde{\kappa}$ | $\kappa/\alpha$ | $5 \times 10^{-11}$ | $\tilde{\kappa}$ | $\kappa/\alpha$ | $3.33 \times 10^{-8}$ |
| $\tilde{\xi}$ | $\xi m_0/\alpha$ | $1.67 \times 10^{-18}$ | $\tilde{\beta}$ | $\beta/(\alpha m_0)$ | $1.33 \times 10^{-8}$ |

Let *m*_0_ be the total initial mass of the system. We scale all concentrations with the initial mass *m*_0_ and time with the typical time associated with the growth parameter *α*. The scalings of the variables and dimensionless parameters are then given by

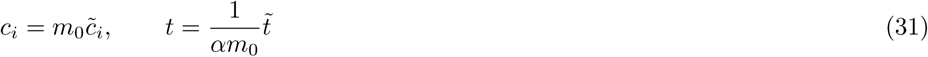

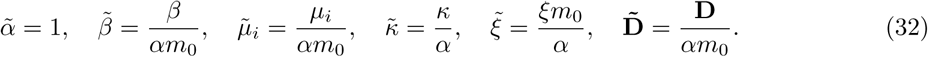

After substitution in the system and then dropping the tildes, we obtain

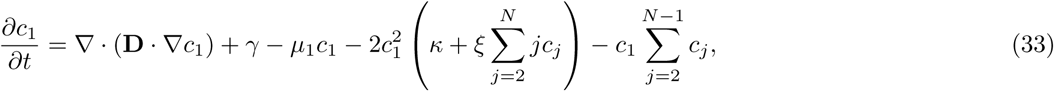

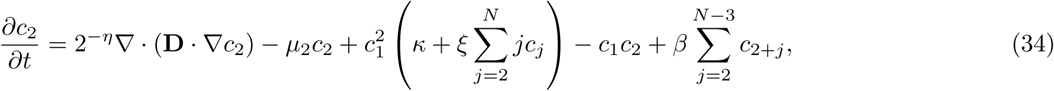

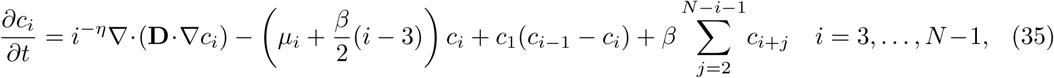

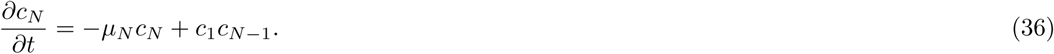

In this new formulation, we have three important small dimensionless parameters (*β* ≪ *κ* ≪ 1 and *ξ, κ* ≪ 1). Note that the rates given in Table 1 are obtained from well-controlled *in vitro* experiments so that they may not be an accurate reflection of the actual process taking place in human brains. Typically, the time scales associated with *in vivo* processes are much longer. In our case, we have typical time scales evolving with dimensionless time *t* ∈ [0, 10^4^]. Nevertheless, they indicate important relative differences in the parameters that we will respect throughout our analysis. For the rest of this paper, we will use the dimensionless version of the equations.

## 4 Smoluchowski network models

It is well appreciated that integrating the continuous equations we have derived for large *N* over the entire brain is extremely difficult even with the most sophisticated methods. By taking advantage of the strong anisotropy of the system, a natural coarse-grained version of the model can be obtained. In this case, we assume that transport only takes place along the axonal pathway and we replace the diffusion operator by the graph Laplacian to obtain a network approximation of the model. These models have been shown to be excellent approximations of the continuous model in the case of the Fischer equation and heterodimer models [51].

For transport along the axon, we model the spreading of monomers and protein aggregates as a diffusion process across the brain’s connectome. The brain connectome is modeled as a weighted graph 𝒢 with *V* nodes (*V* for vertices) and *E* edges obtained from tractography of diffusion tensor images. We can summarize the connectivity of the graph 𝒢 in terms of the weighted adjacency matrix *A*_*ij*_ obtained as the ratio of mean fiber number *n*_*ij*_ and length *l*_*ij*_ between node *i* and *j*. From the weighted adjacency matrix, we compute both the weighted degree matrix *D*_*ii*_, a diagonal matrix that characterizes the degree of each node *i*, and the weighted graph Laplacian *L*_*ij*_ as

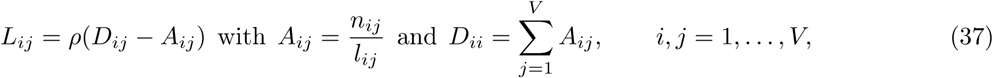

where *ρ* is an overall constant. Since the graph Laplacian has the physical dimension of an inverse length, *ρ* is a velocity and is used to characterize the effective speed at which the diffusion process evolves through the system. We define the concentration *c*_*i,j*_ as the concentration of aggregates of size *i* at node *j*. The particular adjacency matrix that we use for our simulations is obtained from the tractography of diffusion tensor magnetic resonance images of 418 healthy subjects of the Human Connectome Project [52] and is based on the Budapest Reference Connectome v3.0 [53]. The original graph contains 1015 nodes and 37477 edges and it is further reduced here to a graph with *V* = 83 nodes and 1130 edges. The *average path length* (defined as the average number of steps along the shortest paths for all possible pairs of nodes) is 5245*/*3403 ≈ 1.54 and the *global clustering coefficient* (defined as the fraction of paths of length two in the network that are closed over all paths of length two) is 49359*/*69149 ≈ 0.71 which suggests a *small-world network structure*, a fact that has been repeatedly established for brain networks [54]. Further analyses of this network can be found in [51].

The adjacency matrix is shown in Fig. 4 and the graph Laplacian is given explicitly in the Supplementary material as well as the names and positions of each nodes as shown in Fig. 5. For visualization and analysis, we allocate each node to one particular region of the brain, the usual four lobes: *temporal, parietal, frontal occipital*, together with the *basal ganglia*, and the *limbic region*. The last node (shown in black) corresponds to the brain stem.

**Figure 2:**
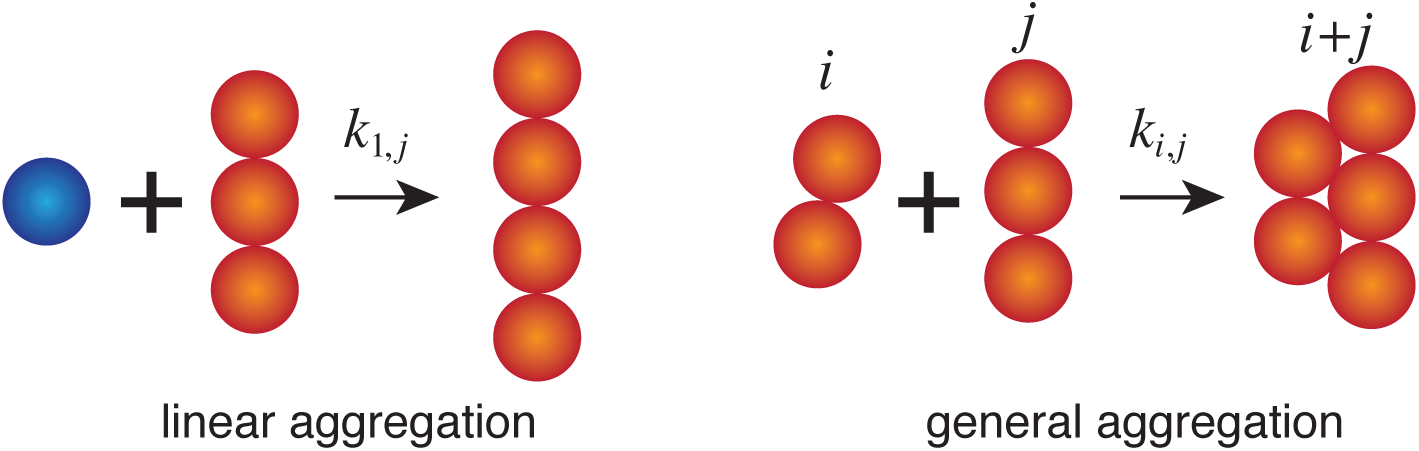
Aggregation processes. During aggregation an *i*-mer merges with a *j*-mer to form an (*i*+*j*)-mer with rate *k*_*i,j*_. Linear aggregation is the particular case where monomers are added to an aggregate and is a good model for fibril formation.

**Figure 3:**
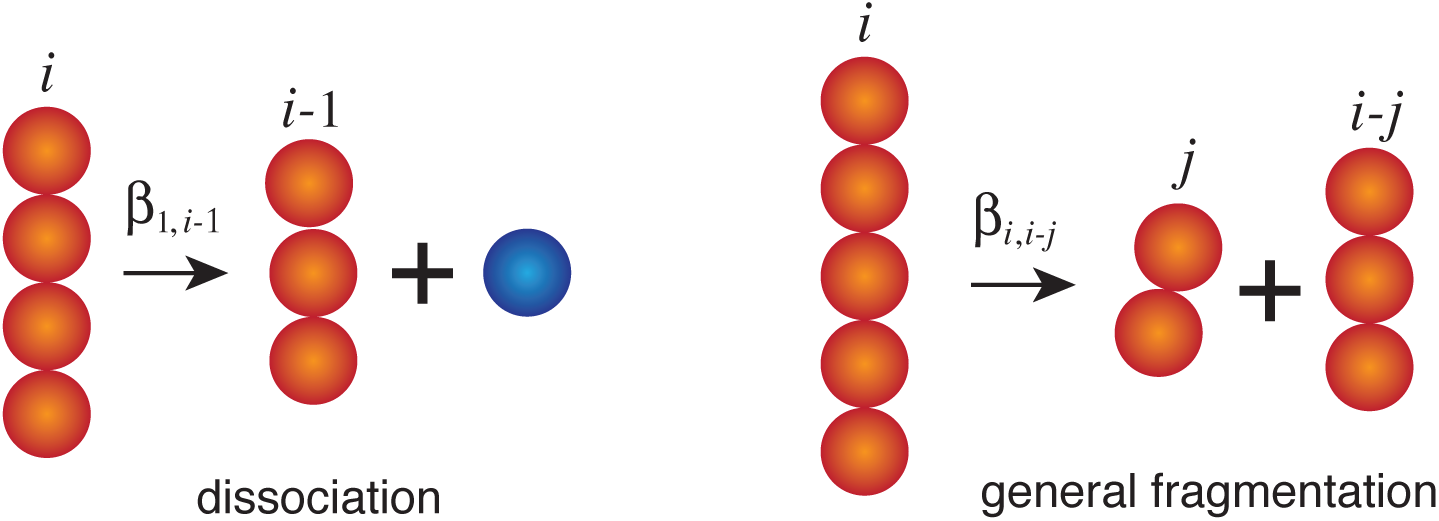
Fragmentation processes. During fragmentation an *i*-mer aggregates with a *j*-mer to form an (*i*−*j*)-mer with rate *β*_*i,i*−*j*_. Dissociation is the particular case where monomers are created.

**Figure 4:**
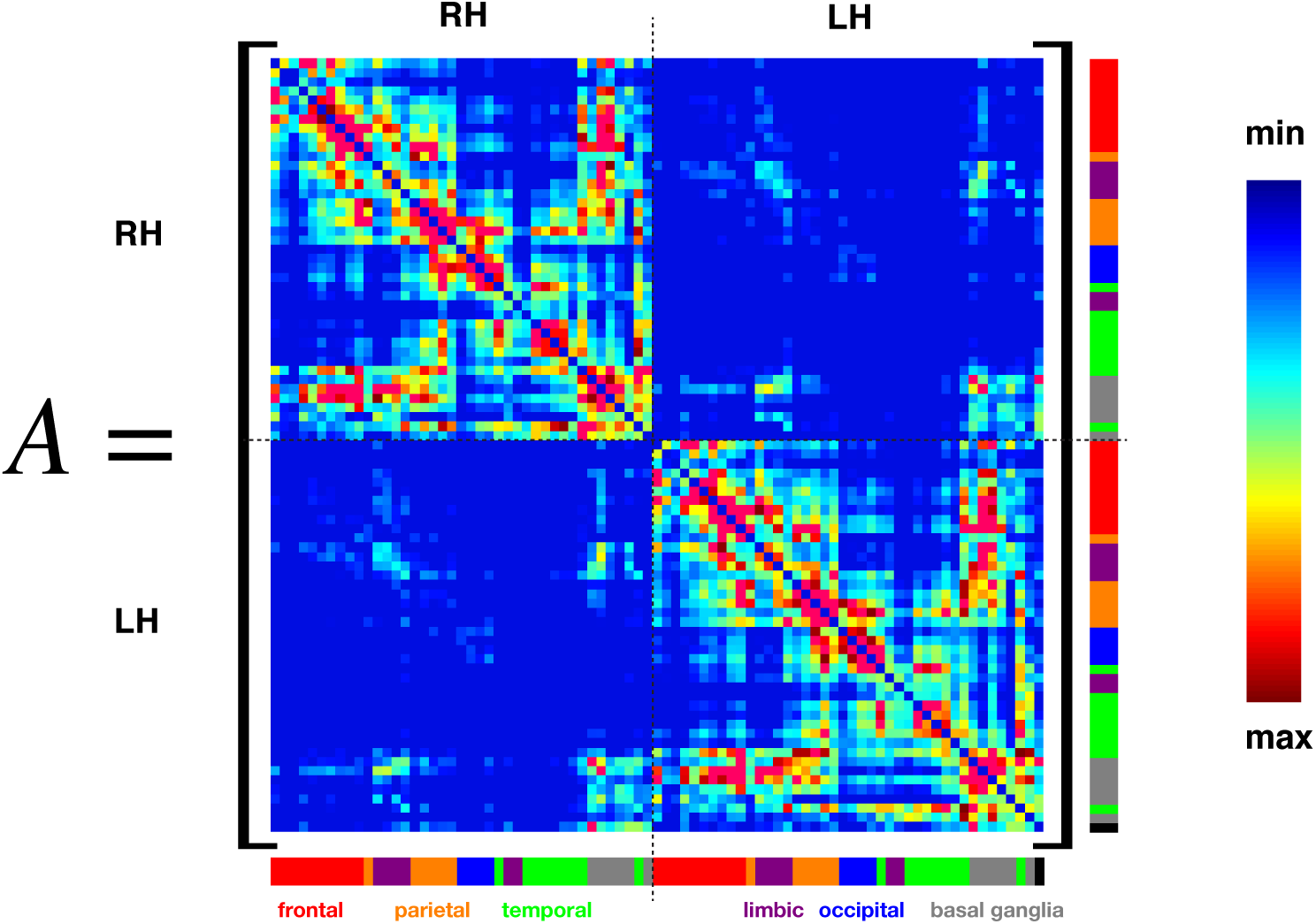
Weighted adjacency matrix with 83 nodes obtained by averaging 418 brains. RH and LH denote the right and left hemisphere, respectively. The color scales from low weight (blue) to high weight (red), the latter indicating strong connections between two nodes. The external color coding around the matrix represents the different regions as depicted in Fig. 5.

**Figure 5:**
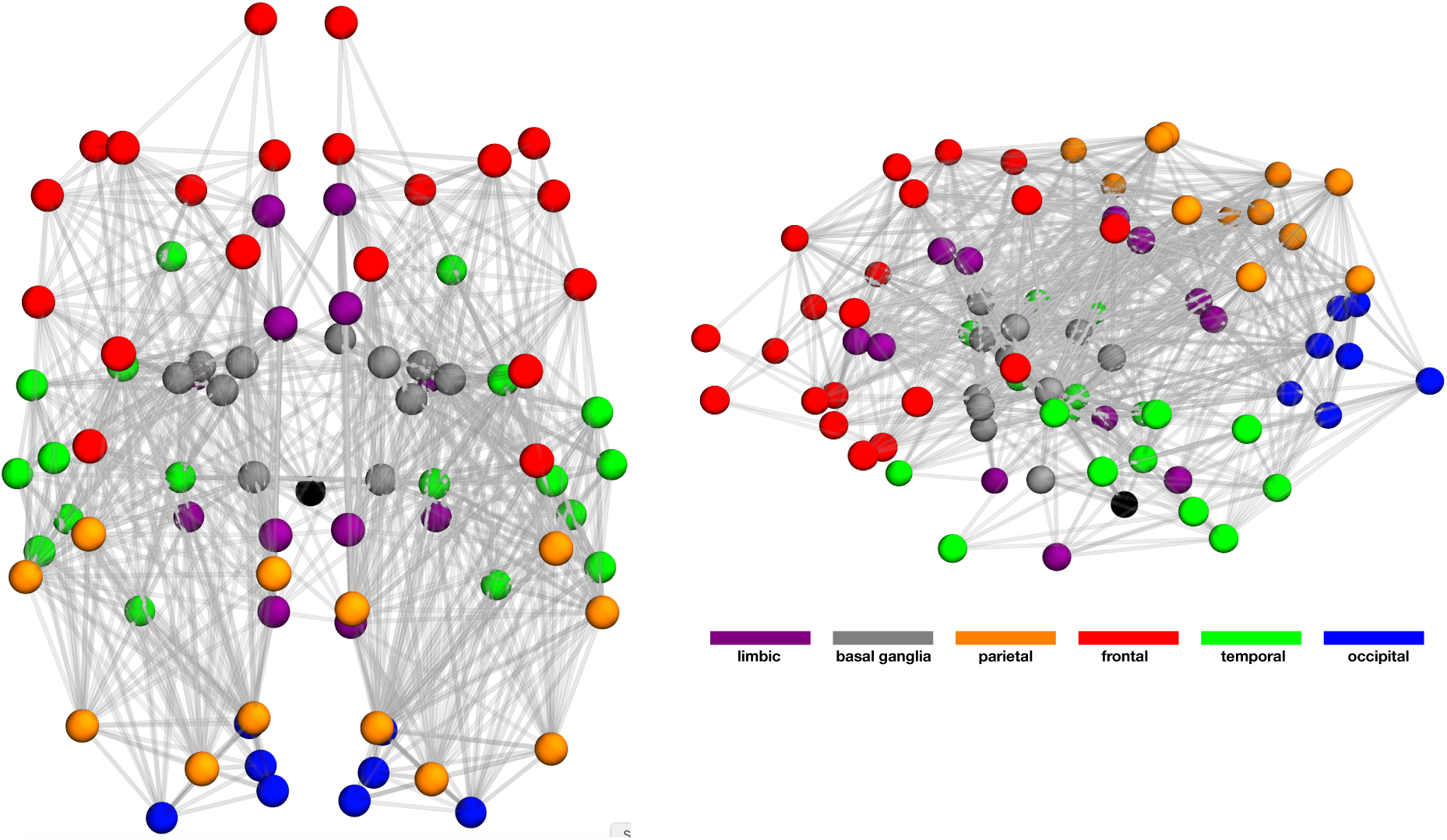
Three-dimensional view of the brain with its six associated regions (the black node denotes the brain stem). Left: view from the top. Right: view from the side.

### 4.1 The network protein model

The network equations corresponding to the continuous model take the form of a system of *N* × *V* first-order ODES:

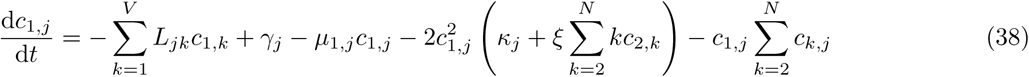

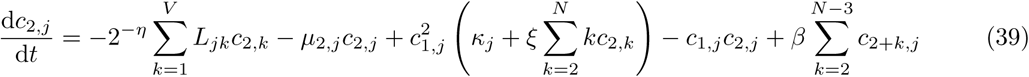

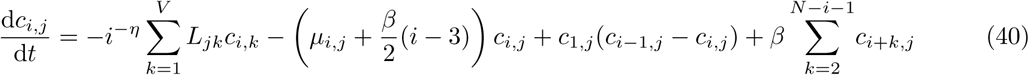

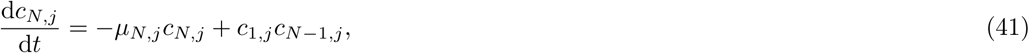

where *i* = 3, …, *N* − 1 and *j* = 1, …, *V*, and we have allowed a possible dependence of the clearance and production rates on the different nodes.

## 5 Analysis of the homogeneous case

### 5.1 Evolution of the total mass

To gain insight into the problem, we start our analysis with the homogeneous case where we look for solutions that are constant in space. In this case, both the network and continuum model lead to same set of ordinary differential equations:

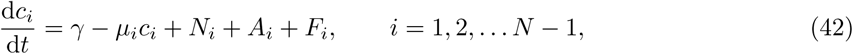

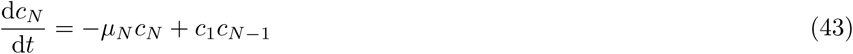

where the different terms take their respective values for the different cases. Two important global quantifiers of the dynamics are the total number of aggregates *P*_tot_ and the total mass *M*_tot_ (or equivalently, total density at constant volume), given by:

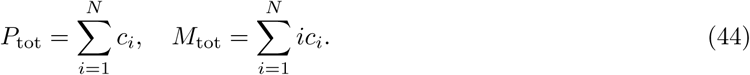

If *N* → ∞, gelation can occur in the system depending on the aggregation law. In this case, mass is not conserved [24]. However, for finite *N*, the evolution of the mass is given by

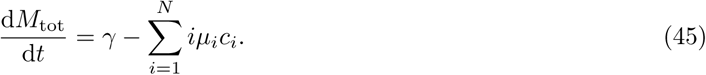

If clearance is size-independent with *µ*_*i*_ = *µ*, then

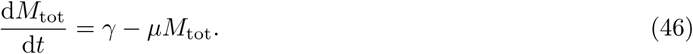

Assuming that at time *t* = 0, *M*_tot_(0) = *c*_1_(0) = 1 = *γ/µ*, then the total mass is conserved and stable (against small perturbation of the initial state). Due to the choice of scaling, we have *M*_tot_ = 1. Starting with an initial population of monomers *c*_1_(0), the total mass remains constant while creating aggregates at the expense of the monomer population. This process does not depend on the particular choice of aggregation process as long as gelation does not take place. In a finite system, gelation is equivalent to treating the super-particle separately. Since there is a finite net flux towards the super-particle, the mass of the other aggregates is lost to the super-particle.

If clearance is size-dependent with *µ*_*i*_ = *µ/i*, then

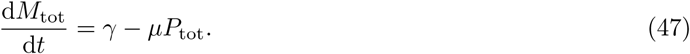

Starting again with *M*_tot_(0) = *c*_1_(0) = 1 = *γ/µ*, we note that since *P*_tot_ ≤ *M*_tot_ and the equality only occurs if *M*_tot_ = *P*_tot_ = *c*_1_, we have 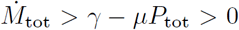 for *t* > 0 and the total mass of the system increases by the creation of new monomers. Particles belonging to aggregates are removed from the system by clearance but their removal is slower than the removal of monomers.

More generally, if we have *µ*_*i*_ ≤ *µ*_1_, ∀*i* > 1 and there is at least one *k* > 1 such that *µ*_*k*_ < *µ*_1_, then, following the same reasoning and initial condition, we have again 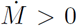 for *t* > 0. The total mass of the system increases.

### 5.2 Moment analysis and evolution of the toxic mass

For the rest of the analysis of the homogeneous system, we will assume that *µ*_*i*_ = *µ* for all *i* and that *N* is sufficiently large as to not affect the dynamics on intermediate time scales of disease progression. Therefore, it is possible to study the limit *N* → ∞. Further, we are interested in solutions with no initial seeding, so that *c*_*i*_(0) = 0, *i* > 1 and *c*_1_(0) = 1. As the system involves, we have, for all time, *c*_1_(*t*) ∈ [0, 1] and *M* (*t*) ∈ [0, 1]. This choice of initial conditions also implies that *γ* = *µ*. The homogeneous system now reads

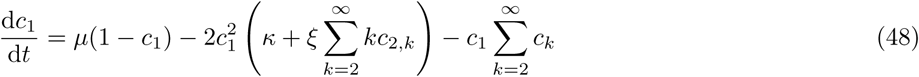

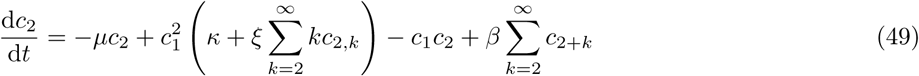

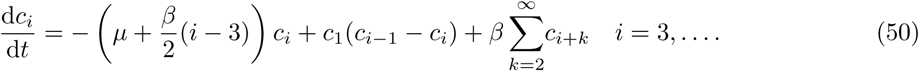

### 5.3 Moment analysis and evolution of the toxic mass

A classic approach to study the infinite system of ODEs (48-50) is to obtain equations for the moments

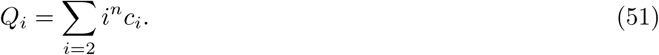

In particular, the first two moments are associated with the total number of toxic aggregates *P* = *Q*_0_ and the total mass *M* = *Q*_1_, respectively. We note that the third moment *Q*_2_ does not appear in this description. This is due to the fact that the two terms involving *Q*_2_ are −1*/*2Σ*i*(*i*−3)*c*_*i*_ and 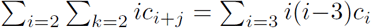, and they cancel exactly. The same cancellation occurs in the model of prion growth [55, 56, 21] and the solution of the resulting closed system can be obtained approximately [57]. This fact has been used by many authors to match experimental data with model prediction [58, 59, 60].

For our model, using the scaled system (48-50), the definition (14), and *m*(*t*) = *c*_1_(*t*), we obtain

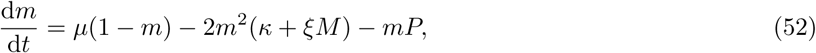

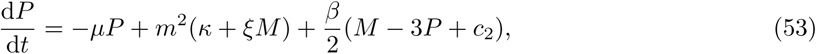

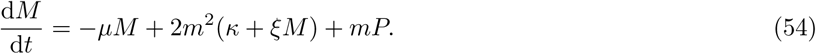

As expected we have *m* + *M* = *M*_tot_ and

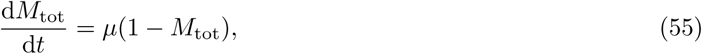

and for the initial condition *m*(0) = 1, *M* (0) = *P* (0) = 0, we have *m*(*t*) = 1 − *M* (*t*) ∀*t*. We note that the system (52-54) is not closed when *β* ≠ 0 as it contains the variable *c*_2_.

### 5.4 Analysis of the A*β* model

Taking *β* = 0 in the moment equations (52–54) leads to a closed system for (*m, P, M*):

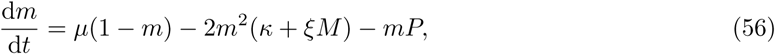

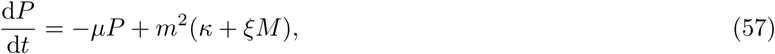

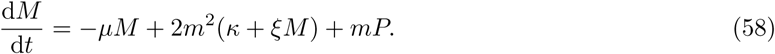

As shown in Fig. 6, uts dynamics from the initial condition (1, 0, 0) tends asymptotically to a fixed point (*m*_∞_, *P*_∞_, *M*_∞_) where *m*_∞_ is the first positive root of

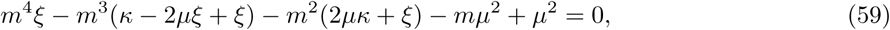

and

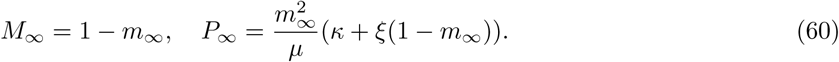

From the asymptotic values, we can determine the exact asymptotic distribution by finding the equilibria of (48-50) in the case *β* = 0:

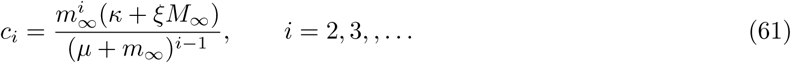

an example of which is shown in Fig. 7 together with a numerical solution of the dynamics leading to the asymptotic distribution. We note that the dynamic is associated with multiple time scales. Initially, the population of toxic protein increases exponentially with a typical time scale obtained by assuming that *m*(*t*) ≈ 1. But, the size distribution only reaches its asymptotic value over a much longer typical time scale compared to the mass of toxic protein. Using *m*(*t*) = 1 in (56-58) leads to a linear system with early time dynamics

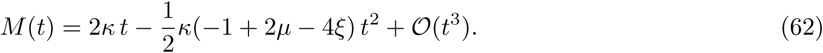

The time at which this solution reaches the asymptotic value *M*_∞_ provides an estimate for the time scale of early expansion of the toxic proteins

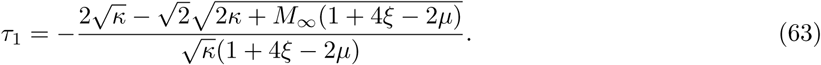

The initial expansion phase is followed by a saturation stage with time scale *τ*_2_ at which *m* is close to its asymptotic value *m*_∞_

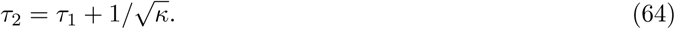

For longer times, the system exhibits a slower dynamical evolution over a time scale *τ*_*n*_ > *τ*_2_ for large *n* towards the asymptotic size distribution. Indeed, once *c*_1_ is closed to its asymptotic value, the equations for *c*_*i*_ with *i* > 2 becomes linear with a typical decay rate given by 1*/*(*µ* + *m*_∞_). Hence the concentration *c*_*n*_ reaches equilibrium on a time scale

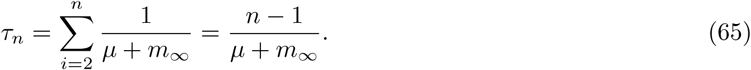

A couple of examples of these time scales are shown in Fig. 7.

**Figure 6:**
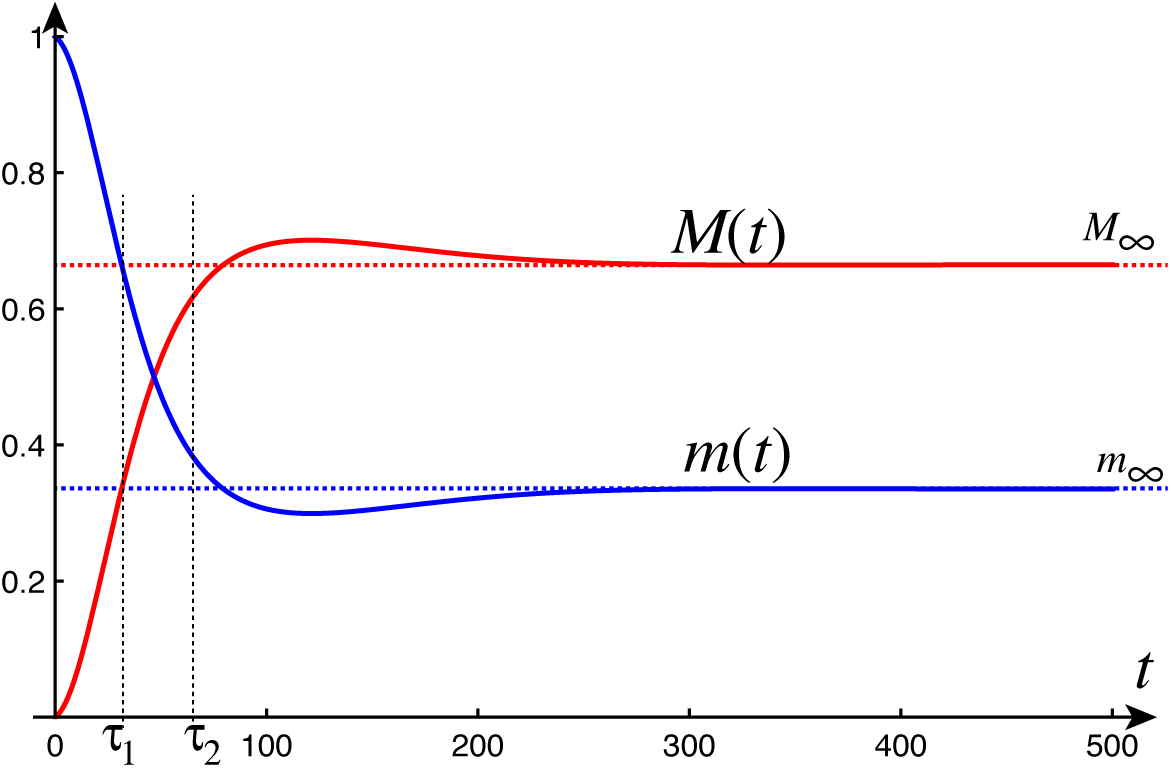
Dynamics of monomer and toxic protein mass in the A*β* model. Here, the initial exponential growth of the toxic population is associated with the time scale *τ*_1_ ≈ 35 and *τ*_2_ ≈66. Parameters: *µ* = 10^−2^, *κ* = *ξ* = 10^−3^.

**Figure 7:**
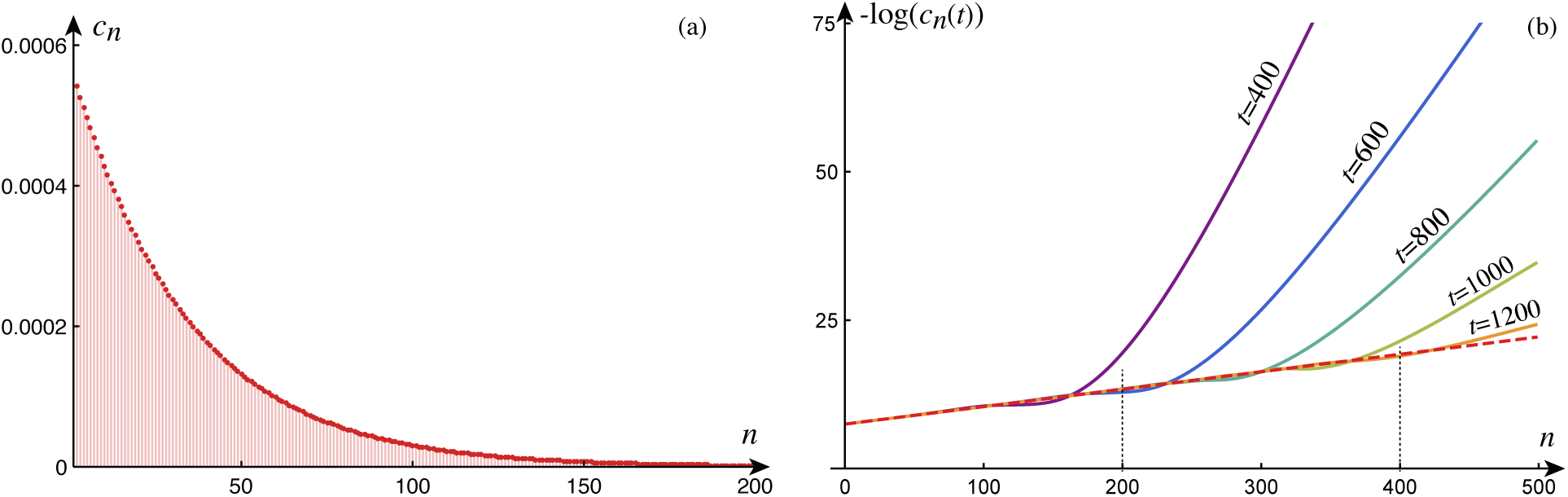
(a) Asymptotic size-distribution in the homogeneous A*β* model. (b) Dynamic evolution of the size distribution (dashed curves indicates the exact asymptotic solution). The typical time scale for an aggregate of size *n* to reach equilibrium is *τ*_*n*_. For instance, here *τ*_200_ ≈ 574 and *τ*_400_ ≈ 1156. Parameters: *µ* = 10^−2^, *κ* = *ξ* = 10^−3^.

### 5.5 Analysis of the *τ* model

The moment equations for the *τ* model read

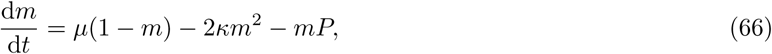

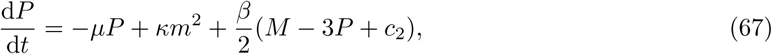

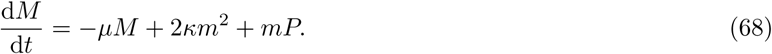

The analysis of these equations is complicated by the fact that they involved *c*_2_(*t*). However, since *β* ≪ 1 and we expect *c*_2_ to be also small, this term can be neglected in the first instance to obtain an approximate but closed system for the moments. Indeed, Fig. 8 shows that the numerical solutions of the full system is indistinguishable from the approximate moment equations. To make this argument more precise, we can obtain exact upper and lower bounds for the asymptotic monomer mass *m*_∞_ by realizing that, asymptotically, *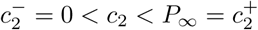*, where

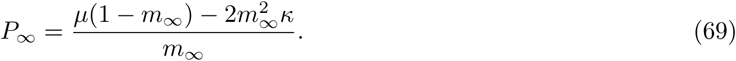

Then, the asymptotic concentration of monomer *m*_∞_ is sandwiched between 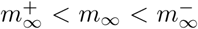 where 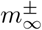 are the two real solutions of

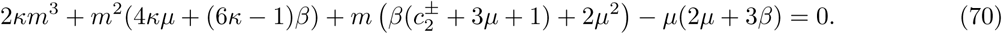

The asymptotic size distribution can be obtained by solving numerically the full system (48-50) for time as shown in Fig. 9. The early dynamics is dominated by the nucleation process with a typical time scale 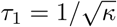. We note that the size distribution is markedly different compared to the A*β* model with a maximum at a value less than the average length given by *M*_∞_*/P*_∞_. In order to obtain an estimate of this asymptotic profile, we assume that we know from the moment equation the asymptotic values of both the monomer population *m*_∞_ and the total aggregate number *P*_∞_ from the previous argument. The problem is then to find a solution for the infinite set of equations

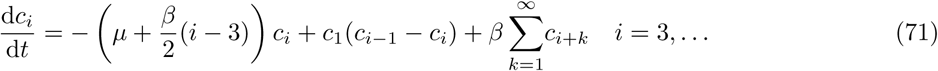

where we have approximated Eq. (50) by changing the summation index in the last term (from *k* = 2 to *k* = 1). We can obtain a continuous limit of this equation by assuming that it is a discretization of an equation for the variable *y*(*s, t*) such that *y*(*n, t*) = *c*_*n*_(*t*). The difference between two consecutive equations of the form (71) is

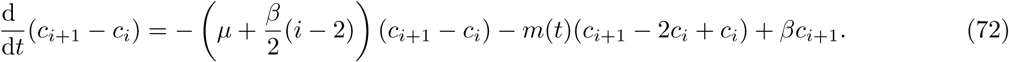

Using the discretization of *y* with a unit step, we have

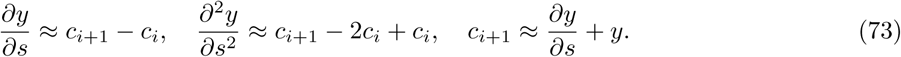

Equation (72) can then be written

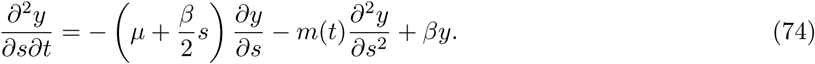

The steady state of this equation is given by the solution of

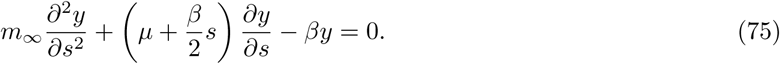

This is a linear second-order equation for *y*. Enforcing that the solution at *s* = 0 is bounded leads to a solution with a single constant:

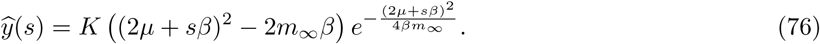

The constant *K* is found by the condition

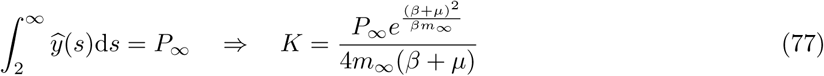

This solution, shown in Fig. 9, is a good approximation of the exact discrete distribution. In particular, it gives an excellent estimate for the maximum, located at the closest integer to

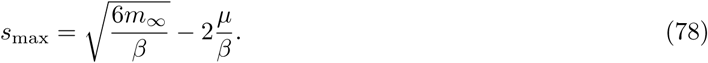

This estimate also shows that fragmentation is necessary to observe a maximum away from *N* = 2.

**Figure 8:**
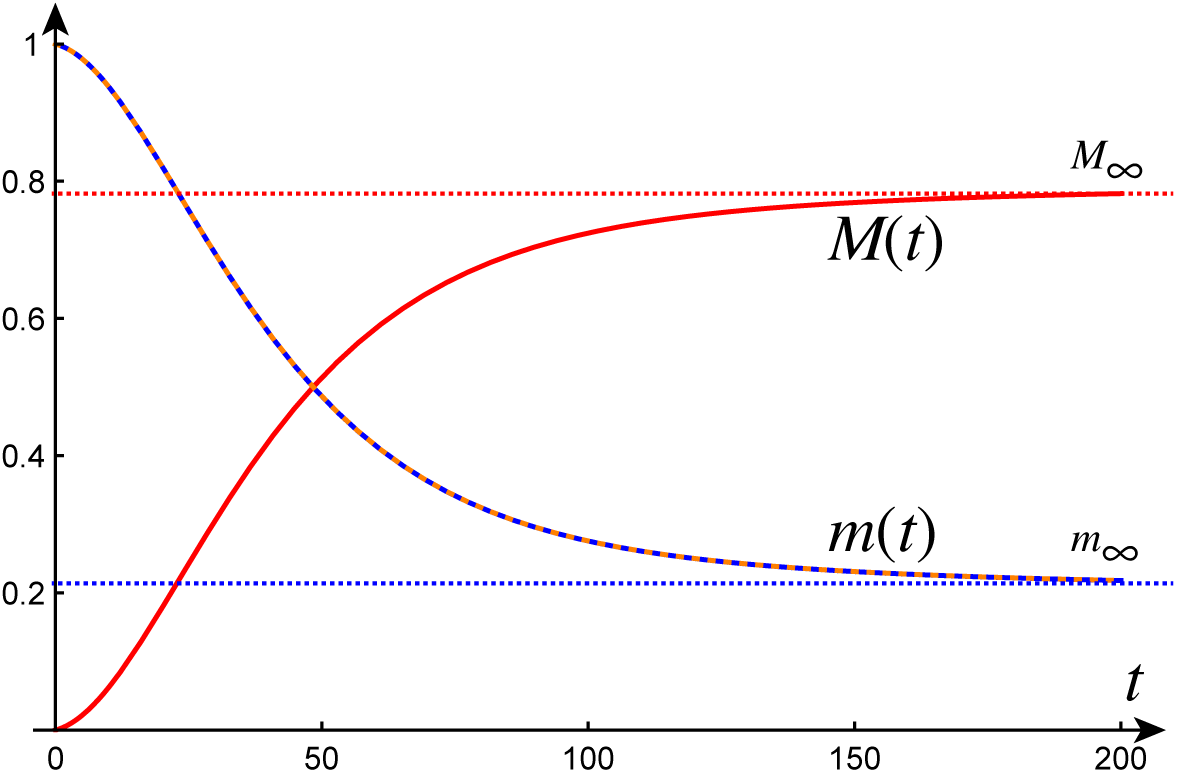
Dynamics of monomer and toxic protein mass in the *τ* model. The dashed line indicates the solution of the moment equations for the monomers obtained by setting *c*_2_ = 0. Parameters: *µ* = 10^−2^, *κ* = *β* = 10^−3^.

**Figure 9:**
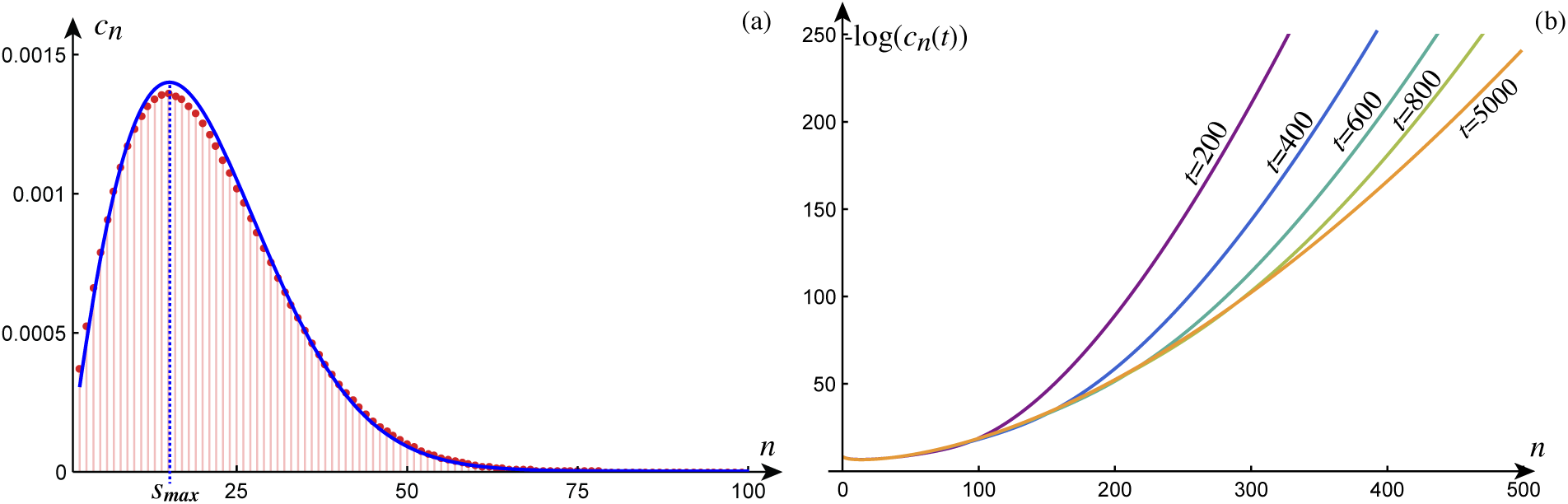
(a) Asymptotic size-distribution in the homogeneous *τ* model. Here, the average length given is *M*_∞_*/P*_∞_ ≈ 21 and the maximum is reached at size 15. The continuous (blue) curve is the continuum approximation. The predicted maximum also occurs at 15, the closest integer to *s*_max_ ≈ 15.12. (b) Dynamic evolution of the size distribution. Parameters: *µ* = 10^−2^, *κ* = *β* = 10^−3^.

The asymptotic dynamics for large *t* can be found by analyzing (74) and looking for solutions of the form

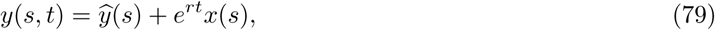

which leads to an equation of the form (75) where *y* is replaced by *x* and *m*_∞_ is replaced by *m*_∞_ + *r*. This equation has two solutions and the conditions that this time-dependent solution preserves both the number and mass of aggregates

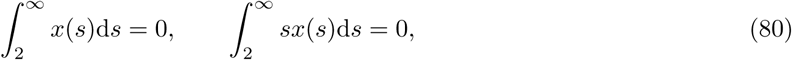

lead to an equation for *r*

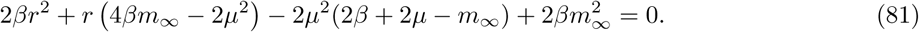

Both solutions are valid but the largest negative solution is the solution of interest for the dynamics. Indeed the smallest negative exponent describes solutions that quickly decay to the static solution. The solution associated with the largest exponent is the one observed for large times.

entorhinal

## 6 Network simulation

Next, we consider the dynamic evolution of protein concentrations at the level of the network. The first question is to scale parameters and variable correctly from the homogeneous equations studied in the previous section and valid at one node to the entire network. The total mass of monomer in the system *m*_0_ is assumed to be distributed uniformly on all the nodes so that, in the scaled variables (31), the initial conditions for the network are

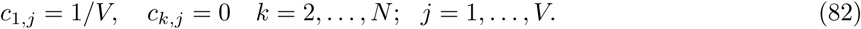

Then, for the network to have the same kinetics as the homogeneous system, we must scale the parameters from the homogenous system (now described by the subscripts “hom”) as follows

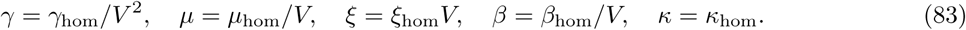

Similarly the time scale is now *t* = *t*_hom_*V*. The equivalence with the homogenous system is obtained by setting *κ*_*i*_ = *κ* and *ρ* = 0 in (38-41). Then the total mass of monomers *m* = Σ*c*_1,*j*_ and toxic proteins 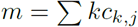 has the same dynamics as the one obtained in Fig. 6 and 8.

The second question is how to properly seed the system to express the fact that the diseases start at a given location. We can either start with a non-zero initial condition of dimers at a given node or, assume that the main mechanism for the initial creation of toxic proteins is due to nucleation at a given node. Here, we choose the latter modeling assumption and assume that *κ*_*i*_ vanishes everywhere except at given nodes where it assumes a small value. These nodes are the seeding regions were neurodegenerative diseases are known to start. For the A*β* model we start at the two nodes characterizing the posterior cingulate [61]. For the *τ* model, we seed the system in the entorhinal region [62] (the list of regions of interest, together with their node number, lobe, hemisphere, and spatial coordinates is given in the Supplementary Material).

Since the diffusion tends to homogenize the system, we expect that for long times the dynamics is uniform over all the nodes so that the size distribution is described by the homogeneous system.

From now on, we assume that the clearance rate for each aggregate is the same so that the total mass of proteins is conserved.

### 6.1 Comparison of the A*β* and *τ* models

For both systems, we use the same parameters apart for the fragmentation (*β* = 0 for the A*β* model), the secondary nucleation (*ξ* = 0 for the *τ* model) and the seeding region as described above. For the simulations we chose the parameters given in Table 2. Note that the asymptotic decay in size is faster with the *τ* model. Hence, for the values of the parameters chosen here, we only need to consider aggregates of size up to *N* = 200 as the concentration of large aggregates is negligible.

**Table 2:** Parameters chosen for the numerical simulation based on the analysis of the homogeneous system (c. stands for concentration).

| $A\beta$ model | | | $\tau$ model | | |
| --- | --- | --- | --- | --- | --- |
| $\kappa_i$ | nucleation at node $i$ | $10^{-3}(\delta_{i,14} + \delta_{i,55})$ | $\kappa_i$ | nucleation at node $i$ | $10^{-3}(\delta_{i,27} + \delta_{i,68})$ |
| $\xi$ | secondary nucleation | $10^{-3}V$ | $\xi$ | secondary nucleation | 0 |
| $\beta$ | fragmentation rate | 0 | $\beta$ | fragmentation rate | $10^{-3}/V$ |
| $\alpha$ | elongation rate | 1 | $\alpha$ | elongation rate | 1 |
| $\gamma$ | production rate | $10^{-2}/V^2$ | $\gamma$ | production rate | $10^{-2}/V^2$ |
| $\mu$ | clearance rate | $10^{-2}/V$ | $\mu$ | clearance rate | $10^{-2}/V$ |
| $\rho$ | diffusion constant | $2 \times 10^{-5}$ | $\rho$ | diffusion constant | $2 \times 10^{-5}$ |
| $m_0$ | initial monomer c. | 1 | $m_0$ | initial monomer c. | 1 |
| $V$ | number of nodes | 83 | $V$ | number of nodes | 83 |
| $N$ | super-particle size | 400 | $N$ | super-particle size | 200 |

#### 6.1.1 Evolution of total monomer and toxic protein mass

Despite the fact that the system is not homogeneous, the overall total toxic mass (obtained by summing the mass of each aggregates at each node) follows a similar evolution as the homogeneous system. For the A*β* model, the asymptotic value for *m*_∞_ can be obtained by using (59) for the entire system after the proper scaling of the parameters

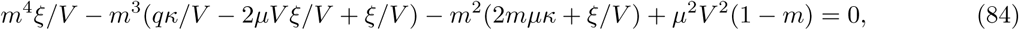

where *q* is the number of nodes seeded (2 in our case). For the *τ* model, we use the network version of (70)

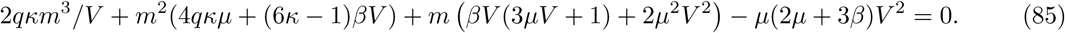

The evolution for the particular choice of parameters in Table 2 is shown in Fig. 10.

**Figure 10:**
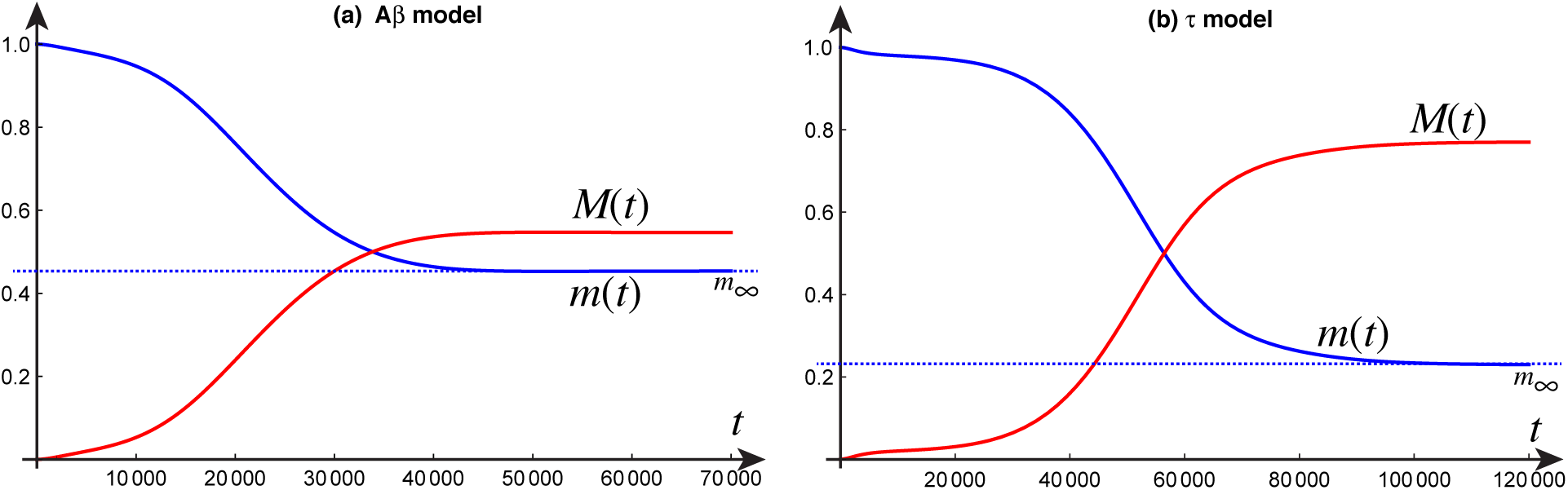
The total monomer concentration *m*(*t*) (the sum of *c*_1,*i*_ over all the nodes) and the total mass of aggregates *M* (*t*) = 1 − *m*(*t*) for (a) the A*β* model (with both estimated and numerical asymptotic value given by *m*_∞_ ≈ 0.45); and (b) the *τ* model (with both estimated and numerical asymptotic value given by *m*_∞_ ≈ 0.23). Parameters given in Table 2.

#### 6.1.2 Evolution of the size distribution

We compute for the values given in Table 2, the evolution of the size distribution for both models. We can obtain asymptotic estimates based on the same argument as in the previous section. For the A*β* model, we have

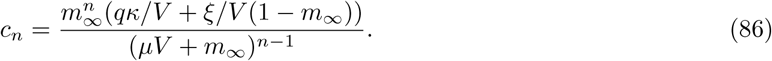

For the *τ* model, we have

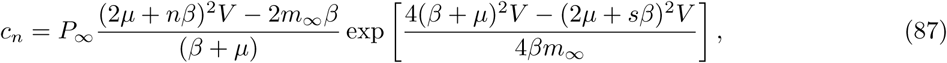

with a maximum at the closest integer to

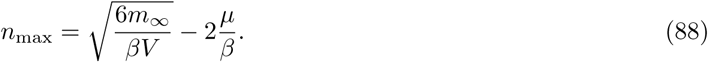

We notice that the two models exhibit distinct size distribution and that for the *τ* model, long chains (of size > 40 monomers) tends to disappear quickly in the long-time dynamics, while this is not the case for the *Aβ* model that has a much longer-tailed distribution.

#### 6.1.3 Spreading behavior over the network

To understand the evolution of the toxic proteins over the entire network, we compute at each node the toxic mass as a function of time:

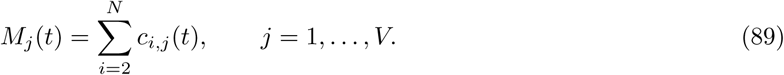

We also average the toxic mass for six regions (consisting of the usual four lobes: *temporal, parietal, frontal occipital*, together with the *basal ganglia*, and the *limbic region* shown in Fig. 5):

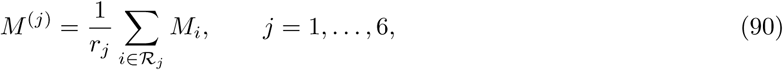

Where ℛ_*j*_ is defined as the set of all nodes in that region and *r*_*j*_ is the number of elements of ℛ_*j*_.

The evolution of the toxic mass at each node clearly illustrated the extra delay in the spreading of the disease associated with diffusion from one node to the next. While the progression at the seeding node is very fast, other nodes feel the effect of the disease over a new time scale directly associated with diffusion (through the overall scaling constant *ρ*). The last node to be invaded is the frontal pole sitting at the extremity of the frontal lobe and poorly connected in the connectome. If these extreme nodes are removed from the computation, the occipital lobe becomes the last lobe to be fully infected.

#### 6.1.4 Staging estimates

A striking features of Fig. 12ab is that staging is established very early on in the dynamics. Once the process starts the ordering of nodes by the toxic mass does not change significantly (no curves intersect).

This observation can be used to provide an estimate of the spatial staging. Indeed, for early times, the only significant change in the system is a conversion from a large population of healthy monomers to dimers. Therefore, we have *c*_1,*j*_ ≈ 1*/V* and *c*_*i,j*_ ≈ 0 for all *i* > 2. Denoting *q*_*j*_ = *c*_2,*j*_ and using Eq. (39), the dynamics of dimers for early time is therefore approximated by

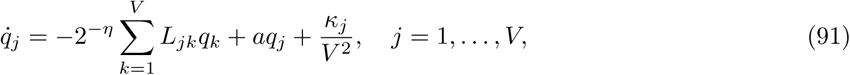

where *a* = 2*ξ/V*^2^ − 1*/V µ*_2_. This is a linear system of ODEs with constant coefficients that can be solved using traditional methods of diagonalization [63, 64]. Indeed, from the graph Laplacian *L*, we can build the matrix *U* whose columns are eigenvectors associated with the eigenvalues of *L*. Introducing the diagonal matrix Λ = diag(*λ*_1_, …, *λ*_*n*_), we have *LU* = *U* Λ. Then, the solution is simply

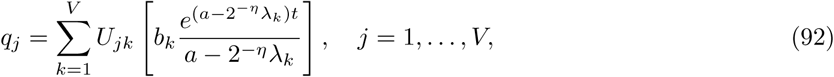

where

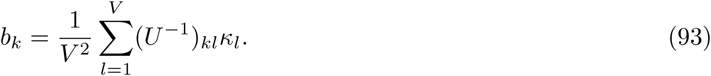

This approximation can be used to sort out the nodes according to the strength of the infection. It provides an excellent overall approximation of the staging that recovers, without the need of any numerical simulation, the overall lobe staging shown in Fig. 12cd. It can also be used to obtain an understanding of the infection process based on the topological properties of the network. Indeed. Expanding this expression for small times, we obtain

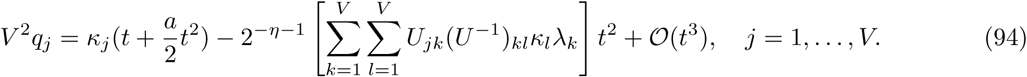

This solution can be further simplified by using *L* = *U* Λ*U*^−1^:

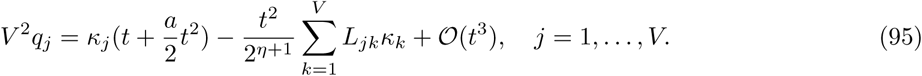

An interesting structure now appears in this solution. We define **q** = (*q*_1_, …, *q*_*V*_), and similarly ***κ*** = (*κ*_1_, …, *κ*_*V*_). Then, after basic alegbraic manipulations, the expansion for the solution **q** can be written

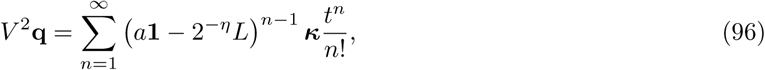

**Figure 11:**
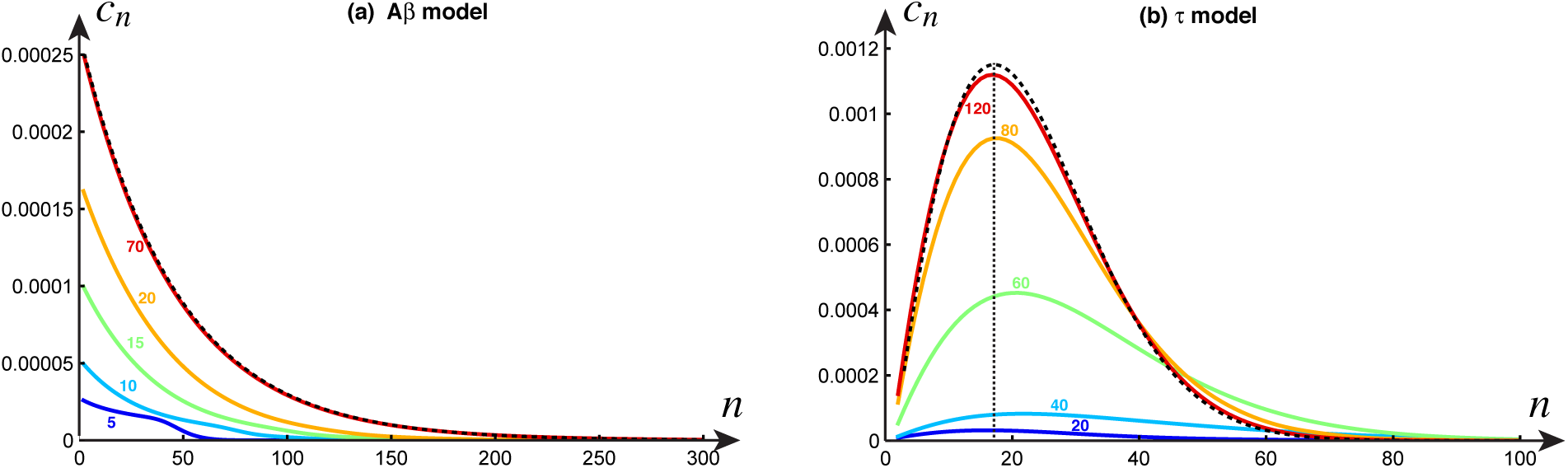
Size distribution for (a) the A*β* model at times 5,000 to 70,000 and (b) the *τ* model at times 20,000 to 120,000 (with estimated and computed *n*_max_ ≈ 17). The dashed lines is the estimated asymptotic distribution. Parameters given in Table 2.

where **1** is the identity matrix.

**Figure 12:**
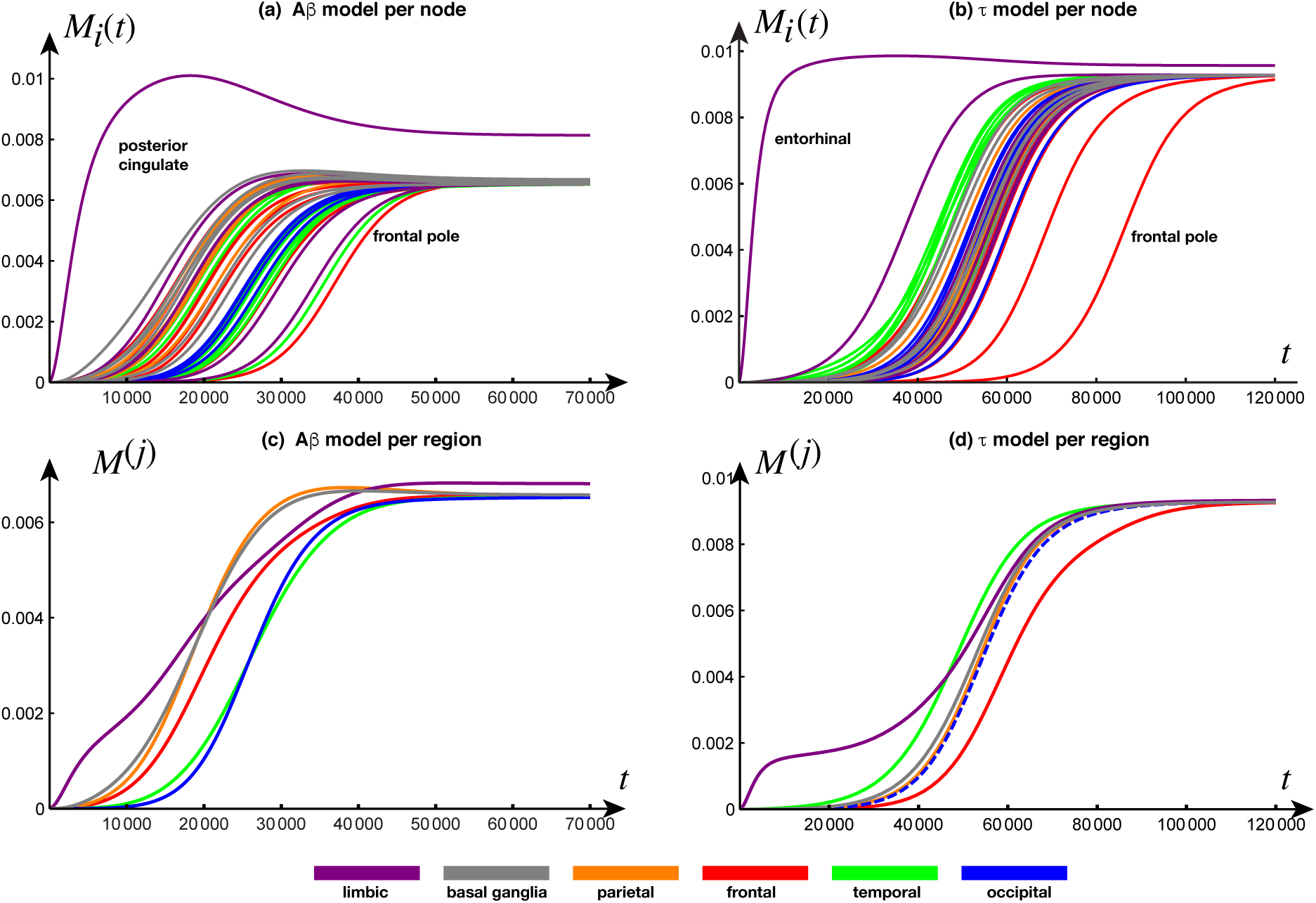
Spreading in the right hemisphere. Top row: Toxic mass at each node in the right hemisphere for (a) the A*β* model and (b) the *τ* model. Bottom row: For each region in the right hemisphere, we show the average toxic mass for (c) the A*β* model and (d) the *τ* model. Parameters given in Table 2.

This expression shows that at very early times, to order 𝒪(*t*), the only nodes affected are the nodes that are seeded (the nodes *j* for which *κ*_*j*_ ≠ 0). This behavior is observed in both Fig. 12ab where the concentration at the seed is seen to increase linearly before it affects other nodes. Later on, to order 𝒪(*t*^2^), the toxic mass increases at the seeded node depending on the kinetics (encoded in the parameter *a*). It also increases at other nodes depending on the product *L****κ***. This product is identically zero for all nodes unless connected to the seeding node. To order 𝒪(*t*^2^), a node *k* has non-zero toxic mass if and only if *L*_*jk*_ ≠ 0 where *j* is one of the seeded nodes. Remarkably this expression mostly depends on the topology of the network (encoded in the matrix *L*). To order 𝒪(*t*^3^), a node has non-zero toxic mass if and only if its path length (the smallest number of steps between two nodes) to a seeded node is two. Hence, to 𝒪(*t*^3^) a new node is seeded only if it is connected to a neighbor of a seeded node. In general, to order 𝒪(*t*^*n*^) a node has non-zero toxic mass if and only if its path length to a seeded node is less than *n*. For instance, the early dynamics of a node *j* that is located at a path length of 5 from a single seeding node *k* will be

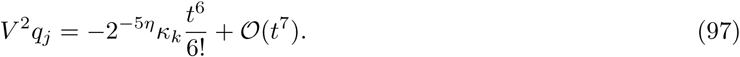

However, due to the small-world structure of the brain network, the average path length is 1.5. Therefore, most nodes connected to the neighbors of the seeded node will have a dynamics starting to order 𝒪(*t*^3^). We conclude that within this model, the following staging dynamics, illustrated in Fig. 13 and Fig. 14 (full movies given in the Supplementary Material), naturally emerges:

- Initial stage: occurring at times 𝒪(*t*) at the seeding nodes (the nodes *j* such that *κ*_*j*_ ≠ 0).
- Primary infection: occurring at times 𝒪(*t*^2^) at the nodes directly connected to the seeding nodes.
- Secondary infection: occurring at times 𝒪(*t*^3^) and depending both on the network topology and the protein kinetics. It affects only regions close to the nodes connected to the seeding nodes.
- Late stage: After the secondary infection, a rapid progression towards an invasion of the entire system takes place. Only nodes that are poorly connected (such as the frontal poles) remain unaffected.

**Figure 13:**
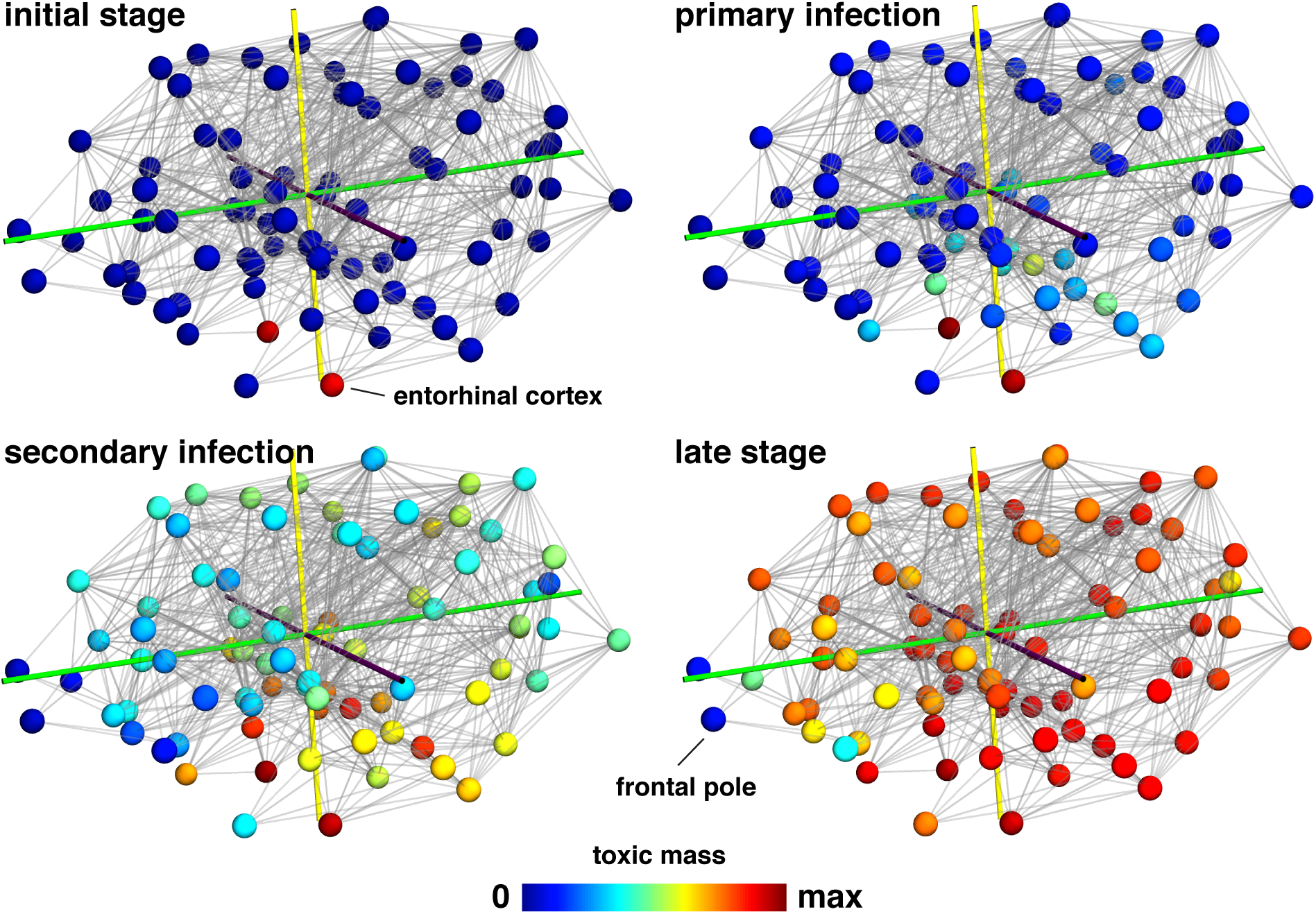
Spatial evolution for *τ* propagation at four time steps corresponding to the initial stage (*t* = 5, 187), primary infection (*t* = 16, 018), secondary infection (*t* = 25, 263), and late stage (*t* = 37, 500). The value of **max** is defined as the maximum value of *M*_*i*_ over all nodes and for all times.

**Figure 14:**
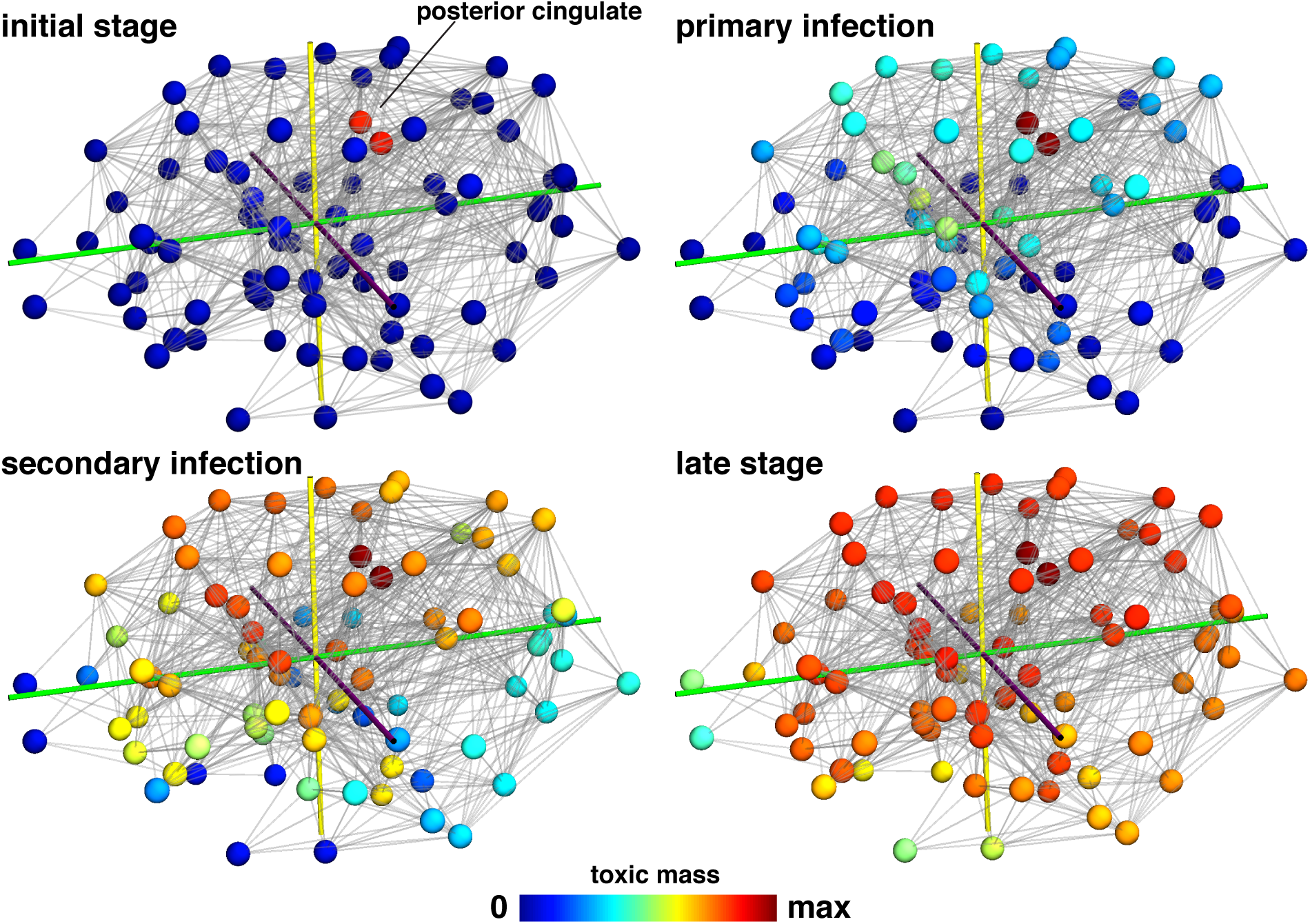
Spatial evolution for A*β* propagation at four time steps corresponding to the initial stage (*t* = 13, 722), primary infection (*t* = 37, 820), secondary infection (*t* = 51, 681), and late stage (*t* = 66, 599). The value of **max** is defined as the maximum value of *M*_*i*_ over all nodes and for all times.

## 7 Conclusion

We have derived a class of models for the spatial progression of key molecules associated with neurodegenerative diseases. These models follow the evolution of aggregates of different sizes and take the form of sets of nonlinear reaction-diffusion equation when the evolution of the aggregates are considered in a continuum with a diffusion evolution along axonal pathways. Taking into account the strong transport anisotropy present in the brain, these equations can be further reduced to systems of Smoluchowski equations interacting on a network through the graph Laplacian. The study of such systems is guided by the homogeneous case for which both total mass evolution and the distribution of aggregates can either be obtained exactly or approximated through various methods by taking advantage of the specific structure of the system.

Here, we considered two paradigmatic cases: a model for the extracellular dispersal of amyloid-*β* further expanded by secondary seeding and aggregation, and a model for the intracellular propagation of *τ* molecules where fragmentation plays a key role. The comparative study of these models shows that fragmentation is key to observe a non-monotonic distribution of aggregate concentrations. It also shows the key role of primary and secondary nucleation processes for the expansion of the initial population.

At the local level, the increase in toxic mass is characterized by an early phase of *seeding* depending on primary nucleation, followed by a period of linear *growth* mostly controlled by aggregation of monomers onto the fibril. Following this early phase, an *expansion phase* takes places that requires secondary nucleation and/or fragmentation. However, if mass is conserved, the expansion phase terminates with a *saturation phase* where toxic and healthy molecules are in balance.

At the global level, the local dynamics is coupled with transport. For a network, the weighted graph Laplacian obtained from tractography provides a direct way to model the transport of toxic aggregates along axonal pathways. The study of the full system for both the A*β* and *τ* models reveals the existence of four different stages in the progression of the disease as shown in Fig. 15. An *initial stage* develops at the seeded node. The evolution is mostly local in time and well described by the homogeneous equation. The *primary infection* takes place in nodes connected to the seeded nodes and mostly depends on the diffusion process rather than the aggregation kinetics. These nodes becomes new seeds and *secondary infection* in all nodes connected to the primary nodes takes place and so on. Recalling that the average path length is about 1.5, it is clear that this network structure leads to rapid infection at this stage of most nodes. In the *late stage*, the disease has invaded all nodes and the toxic mass quickly saturates to its maximal values in balance with the population of healthy monomers.

**Figure 15:**
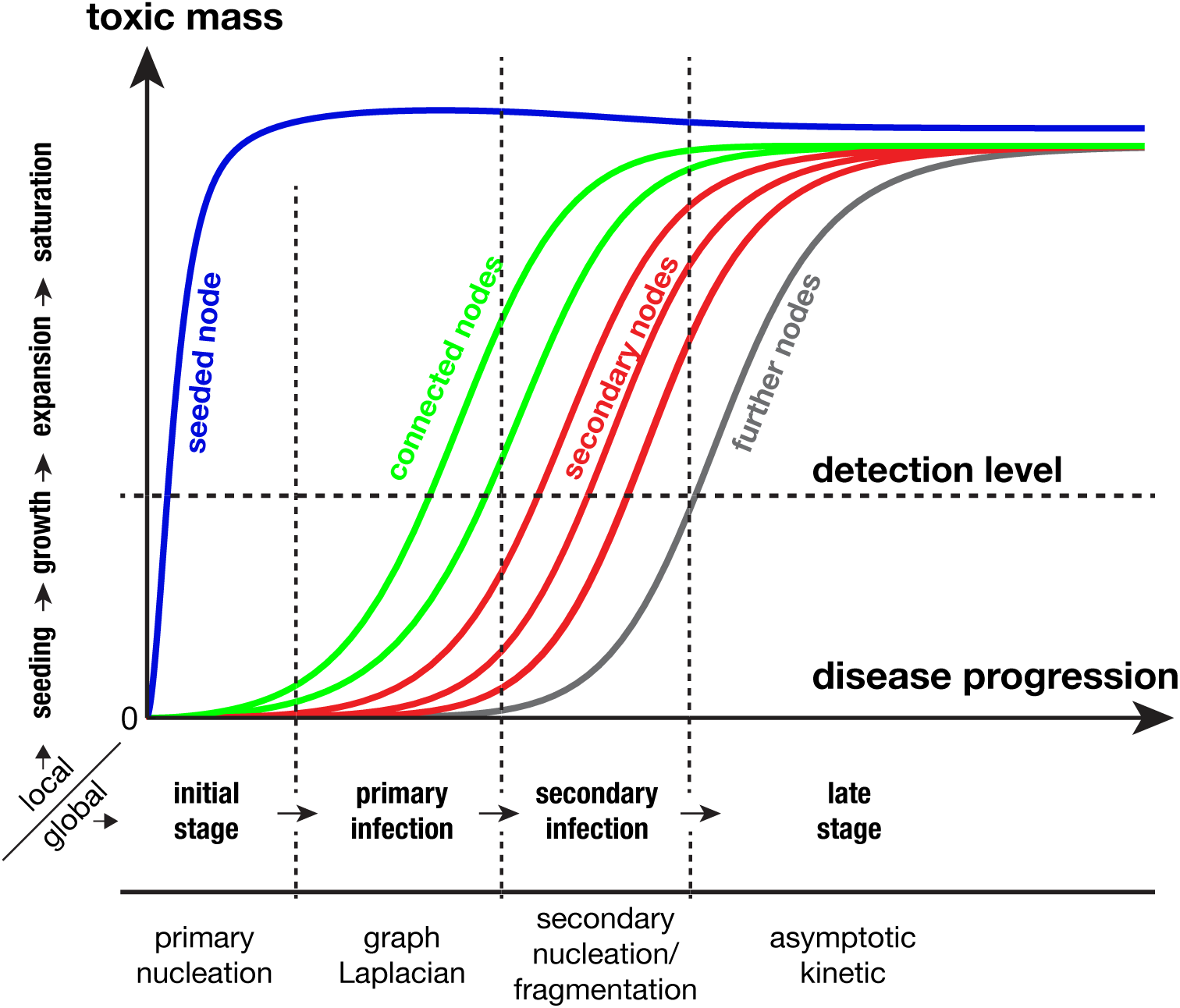
Sketch of the disease evolution as predicted by the model. The local dynamics at each node is shown vertically and starts by seeding, followed by growth, expansion, and saturation. At the global level, the initial stage takes place at the seeded node (blue) and is initially linear in time. Primary infection takes place at nodes (green) connected to the seeded nodes and is driven mostly by the topology of the system as encoded by the graph Laplacian. Secondary infection takes place in secondary nodes (red). These are nodes connected to the primary nodes. The late stage sees a full infection of the system as the concentration of the toxic mass increases both by local kinetics and by diffusion from neighboring nodes. Nodes that are poorly connected to the network are the last affected (gray).

When clearance does not depend on the aggregate size, these models conserve the total initial mass of monomers. This assumption simplifies considerably the study and a complete analysis of the general case, in which clearance varies with aggregate size, would be of great interest. The relative ordering of the parameters we have used in our analysis are based on experimental data. Hence, the typical qualitative features observed in the analysis are universal and directly relevant to the disease progression. Other parameters, such as the effective diffusion constant, production and clearance rates, are not directly accessible based on existing data. The effective diffusion constant in our study has been chosen based on the observation that staging is observed (which implies a very small effective constant). Further study of axonal diffusion in axons and tissues is needed to relate this crucial parameter to microscopic processes. Similarly, it is understood that clearance is key to slow down the progression of the disease. Therefore, a careful analysis of the corresponding parameter and its relationship with other phenomena, such as the vasculature and the glymphatic system, will be crucial in uncovering basic mechanisms and identify possible therapeutic targets.

Theoretical research in neurodegenerative diseases has been so far separated into detailed *in vitro* analysis of aggregation kinetics on the one hand and linear transport on network compared to structural data on the other hand. Both aspects have been shown to be of great importance for our understanding of the diseases. The theory presented here shows that both approaches can be combined within the same mathematical framework and easily analyzed analytically and computationally. The proposed theory is sufficiently flexible to be further generalized to more intricate kinetics or coupled to other important phenomena.

## Supporting information

Full Descrition of SM

## Acknowledgments

This work was supported by the Engineering and Physical Sciences Research Council grant EP/R020205/1 to Alain Goriely and by the National Science Foundation grant CMMI 1727268 to Ellen Kuhl. A.G. gratefully acknowledges a discussion with Tuomas Knowles.

