## Supplementary material for "Spatially-extended nucleation-aggregation-fragmentation models for the dynamics of prion-like neurodegenerative protein-spreading in the brain and its connectome": Full Descrition of SM

The data files are as follows:

GraphLaplacian83.csv is a tab delimited file that contains the weighted graph Laplacian extracted from 418 different brains as explained in the main text.

Names-position-nodes.csv is a tab delimited file that contains 83 rows. Each row has 7 entries: the node number, its hemisphere (left or right), its anatomical name, its coordinates (x,y,z), and its associated region.

3dinvasion-Abeta.mov is a quicktime formatted movie that shows the evolution of toxic mass of amyloid-beta through the connectome

3dinvasion-tau.mov is a quicktime formatted movie that shows the evolution of toxic mass of tau proteins through the connectome
